# Genome-wide and chromosomal microsatellite marker landscape analysis within the genus Crassostrea

**DOI:** 10.1101/2023.12.15.571666

**Authors:** Basanta Pravas Sahu, Mohamed Madhar Fazil, Subhasmita Panda, Vengatesen Thiyagarajan

## Abstract

Microsatellite is a classical codominant marker frequently used to study genetics and evolution of living entities as well as molecular breeding in commercially important species. Although it has a tremendous application in oyster aquaculture, the lack of knowledge about its type, distribution pattern, and comparative analysis is limited. Thus, in this study, we conducted a genome-wide as well as chromosomal microsatellite landscape analysis within the genus Crassostrea. The genome-wide microsatellites number varied from 169432-212368, with relative abundance (RA) and relative density (RD) ranging from 310.18-336.5 loci/Mb and 7553.4-8793.42 bp/Mb, respectively. About 14.99-16.75% of total microsatellites were considered compound microsatellites having cRA and cRD, 21.78-25.5 loci/Mb, and 1332.81-1694.54 bp/Mb, respectively. The mononucleotide microsatellites were predominant followed by di and tetranucleotide. The RA and RD of the SSRs revealed no correlation with genome size but a significant correlation with GC content. However, the number of SSRs showed a significant relationship with the genome size but no relation with GC content. In contrast, the incidence of cSSR was positively associated with genome size and GC content. Finally, 29 cSSR loci were developed and validated in *C. hongkonensis* using one wild population followed by its cross-species amplification. The allele number (*Ne*), observed heterozygosity (*Ho*), expected heterozygosity (*He*), inbreeding co-efficient (*Fis*), the polymorphic information content (*PIC*), ranged from 2-10, 0.092-0.897, 0.0001-1, 0.088-0.828, respectively. The present study elucidated microsatellite evolution within the Crassostrea genome and the loci developed can be utilized for brood stock analysis, parentage assignment, and construction of linkage map of the respective species.

## Introduction

Oysters are bivalve mollusks widely distributed throughout the globe and are considered one of the most economically important seafood (Bailey and Milner 2008). It belongs to the family *Ostreidae* which includes 70 different species of oyster (Sigwart, Wong et al. 2021). Most of them are edible and thus considered an important species for aquaculture (van der Schatte Olivier, Jones et al. 2020). Five commercially important Crassostrea species distributed along the coastal region of China named *C. ariakensis*, *C. gigas*, *C. hongkongensis*, *C. angulata*, and *C. sikamea*. Among them, *C. hongkongensis* is considered a prized oyster species having decent market value with an annual production of more than 1.3 million metric tons (Lam and Morton 2003, Peng, Li et al. 2020). It is an important livelihood of the southern region of China and enriches the mariculture in this area. Recently, seasonal mass mortality and the decrease in quantity and quality of seed have become a matter of concern for this industry (Yu, He et al. 2010, Peng, Liang et al. 2020). Furthermore, over-exploitation and habitat destruction are responsible for the declined population of wild *C. hongkongensis* populations (Caswell, Klein et al. 2020). The lack of knowledge about the genetic diversity that controls the population structure for *C. hongkongenisis* hinders the selection of best quality seeds from the wild for aquaculture farming. Thus, broodstock management and genetic improvement are desirable for the sustainable growth of *C. hongkongensis* aquaculture industry. Although several attempts were made to reveal its genetic diversity using molecular markers (Li, Wu et al. 2013, Ma, Li et al. 2021), still a clear picture is lacking.

Microsatellites, otherwise known as simple sequence repeats (SSRs) composed of 1–6 bp tandem motifs of DNA are prominently observed throughout the eukaryotic and prokaryotic genomes (Tóth, Gáspári et al. 2000, Sahu, Sahoo et al. 2014, Zhang, Chen et al. 2023). Their versatility is characterized by high frequency, codominance, and polymorphic in nature (Vieira, Santini et al. 2016). Furthermore, depending on the composition of their motif, they have been classified as perfect (pSSR/SSR), Imperfect (iSSR), and compound (cSSR) microsatellite (Du, Zhang et al. 2018, Du, Liu et al. 2020). Polymorphic microsatellite markers are frequently used to study genetic variation (Sahoo, Sahu et al. 2014, Chen, Xu et al. 2022); linkage mapping (Sahoo, Patel et al. 2015, Ma, Yu et al. 2022), parentage analysis (Liu, Li et al. 2020), quantitative trait loci analysis (Hornick and Plough 2022).

Comparative genome mapping is one of the important applications for microsatellite mining (Sharma, Grover et al. 2007). Till now, systematic analysis with regard to microsatellite abundance, distribution, differentiation, and characterization have been accomplished on humans (Eckert and Hile 2009), primates (Xu, Li et al. 2018), plants (Kalia, Rai et al. 2011), fungi (Oliveira and Azevedo 2022), viruses (Sahu, Majee et al. 2020, Sahu, Majee et al. 2022), and bacteria (Panda, Swain et al. 2023). By implementing comparative genomics approaches between pigs, cattle, and humans, interspecific microsatellite markers were developed (Sun and Kirkpatrick 1996). A similar study revealed 54% of microsatellite similarity between horse and humans (Farber and Medrano 2004). During the complete genome sequence analysis of large cats, twelve microsatellite loci were identified, which is useful to mitigate the conservation of related species, such as tigers, snow leopards, and lions (Hyun, Pandey et al. 2021). The study of microsatellite presence and its distribution in mammalian genomes has gained much attention, compared with the Ostreidae family. Although several attempts were made to investigate and characterize microsatellite markers for oyster stock and genetic analysis, still a comprehensive analysis is missing (Xiao, Ma et al. 2011, Ma, Zhang et al. 2021, Monteiro, Galvão et al. 2023). Thus, the objectives of this study aims, were 1) genome-wide analysis of abundance and frequency of simple and compound microsatellites within five important oyster genomes, 2) Analysis of microsatellites through chromosomes level and its correlation with length, incidence, abundance, density and GC content, 3) development of cSSR in *C. hongkongenesis* followed by diversity analysis and cross-species amplification.

## 2. Materials and methods

### 2.1. Genomic sequences

Genome sequences from five oyster species, named *C. ariakensis*, *C. gigas*, *C. hongkongensis*, *C. virginica* and *C. angulata* were used in this study as their whole genome as well as chromosomal assembly sequences available in GenBank. Most genome sequences were retrieved from the National Centre for Biotechnology Information (http://www.ncbi.nlm.nih.gov/). The ambiguous nucleotides (Ns) were filtered out by using the Perl script to obtain the valid sequence length. The genome sequences along with their details are listed in Table 1.

**Table 1:** Genome-wide overview of identification and distribution of microsatellites.

| Species | Common Name | Size (MB) | GC % | SSR | RA (loci/Mb) | RD (bp/Mb) | %SSR | Avg length | cSSR | cRA | cRD | cSSR% |
| --- | --- | --- | --- | --- | --- | --- | --- | --- | --- | --- | --- | --- |
| *C. ariakensis* | Suminoe oyster | 663.1 | 33.56 | 206887 | 312.65 | 8238.17 | 0.83 | 26.35 | 14436 | 21.78 | 1490.09 | 14.99 |
| *C. gigas* | Pacific oyster | 647.9 | 33.5 | 194768 | 330.48 | 8793.42 | 0.88 | 26.46 | 16520 | 25.5 | 1694.54 | 16.75 |
| *C. hongkongensis* | Hong Kong oyster | 607.9 | 33.54 | 169432 | 325.14 | 8333.01 | 0.84 | 25.69 | 14998 | 24.64 | 1581.68 | 16.08 |
| *C. virginica* | Eastern oyster | 684.7 | 34.83 | 212368 | 310.18 | 7553.4 | 0.76 | 24.36 | 15175 | 22.16 | 1332.81 | 15.68 |
| *C. angulata* | Portugese oyster | 624.1 | 33.5 | 210075 | 336.5 | 8713.15 | 0.88 | 25.89 | 15744 | 25.22 | 1643.26 | 16.23 |

### 2.2. SSRs identification and its characteristics

The Krait v0.9.0, an ultra-fast and user-friendly software was used for genome-wide investigation of microsatellite markers (Du, Zhang et al. 2018). The parameters for microsatellite investigation set as follows for mono ≥12 bp, di ≥14 bp, tri ≥15 bp, tetra ≥16 bp, penta ≥20 bp, and hexa ≥24 bp, and the optimum length of flanking sequence was ≥ 200 bp, as previously reported (Huang, Li et al. 2016). The maximum distance between two microsatellites (dMax) was 10 bp. Other parameters were set as default to determine real composition. Furthermore, microsatellite prevalence within the genome was deciphered by microsatellite abundance (loci/Mb) and density (bp/Mb) (Qi, Lu et al. 2020).

### 2.3. Primer design for the validation of cSSR

Many studies revealed that cSSR has more potential to be polymorphic compared to simple SSR (Bull, Pabón-Peña et al. 1999). Thus, in this study, we have targeted the compound microsatellite containing nucleotide motifs. Primers were designed by Primer3 with the primer size; 18–27 bp, annealing temperature of the primers (*T*m); 55.0–62.0 °C, GC content; 40–60%, and amplicon product size; 200–500 bp (Sun, Skaletsky et al. 2000).

### 2.4. DNA extraction, PCR amplification, diversity analysis and cross species evaluation

We chose 29 sets of primer pairs for validation of PCR amplification followed by polymorphism analysis. Genomic DNA (gDNA) of 32 individuals of wild *C. hongkongensis* sampled from Hong Kong, Lau Fu Shan province (22.591440;120.333000), used PCR to verify the polymorphism. The amplicons were checked using electrophoresis on 3% agarose gel stained with ethidium bromide. Allele size was analyzed by referring to a 50 bp DNA ladder. Number of alleles (*Ne*), expected heterozygosity (*He*), observed heterozygosity (*Ho*), Hardy-Weinberg equilibrium (*HWE*), Fis inbreeding coefficient (*Fis*), and linkage disequilibrium (*LD*) were determined utilizing GENEPOP 3.4 (Raymond and Rousset 1995). The polymorphism information content (*PIC*) was measured with PIC-CALC v. 0.6. The transferability of all polymorphic cSSR loci developed during this study was evaluated in three closely related species, such as *C. gigas, C. angulata,* and *C. ariakensis*, with five individuals of each collected from Lau Fu Shan Provinces, Hong Kong.

## 3. Results

### 3.1. Genome-wide analysis of the incidence, RA, RD, and GC content of microsatellites

Our study revealed that Crassostrea genomes enriched with SSRs varying from 200631-214550 in number with an average of 208913 per genome. The SSR% within the genome obtained the highest in *C. gigas* (0.88%) and *C. angulata* (0.88%). The RA and RD ranged from 310.01–336.4 loci/Mb and 7550.2–8760.1 bp/Mb, respectively. The lowest RA and RD of microsatellites were found in *C.virginica* (310.01 loci/Mb and 7550.2 bp/Mb). The highest RA and RD were found in *C. angulata* (336.46 loci/Mb) and *C. gigas* (8760.1 bp/Mb), respectively (Table-1, Fig. 1). The SSR% of all five Crassostrea species has the highest frequency of mononucleotide (51.15), followed by di (33.58), tetra (8.82), tri (4.96), penta (0.93) and hexa (0.54) nucleotides (Fig. 2). The GC percentage was obtained for mono to hexanucleotide microsatellites in the five genomes. The overall GC content accounted for approximately 33.5 for *C. areakeensis*, *C. gigas*, *C. hongkongensis*, and *C. angulata* with a maximum of 34.83 in *C. virginica* of the total length of SSRs. The major repeat motifs were (C/G)n, (AC)n, (AT)n, (AG)n, (AAT)n, (AAC)n, (AAAT)n, (AAAC)n, (AAAG)n, (AAGG)n, (AGAT)n and (AAAAC)n shared by the five species. In *C. hongkongensis* (G/C)n mononucleotide repeats were highly abundant as compared to (A/T)n. In contrast, the most frequent dinucleotide repeats are (TC)n, (TA)n, (GA)n, (AT)n, (AG)n, and (CT)n. The trinucleotide repeats were ubiquitously present in *C. hongkongensis* which was composed of (TTA)n (823) followed by (AAT)n (750), (TAT)n (386), (TAA)n (342), and (TTG)n (243) (Fig. 3a, b).

**Figure 1:**
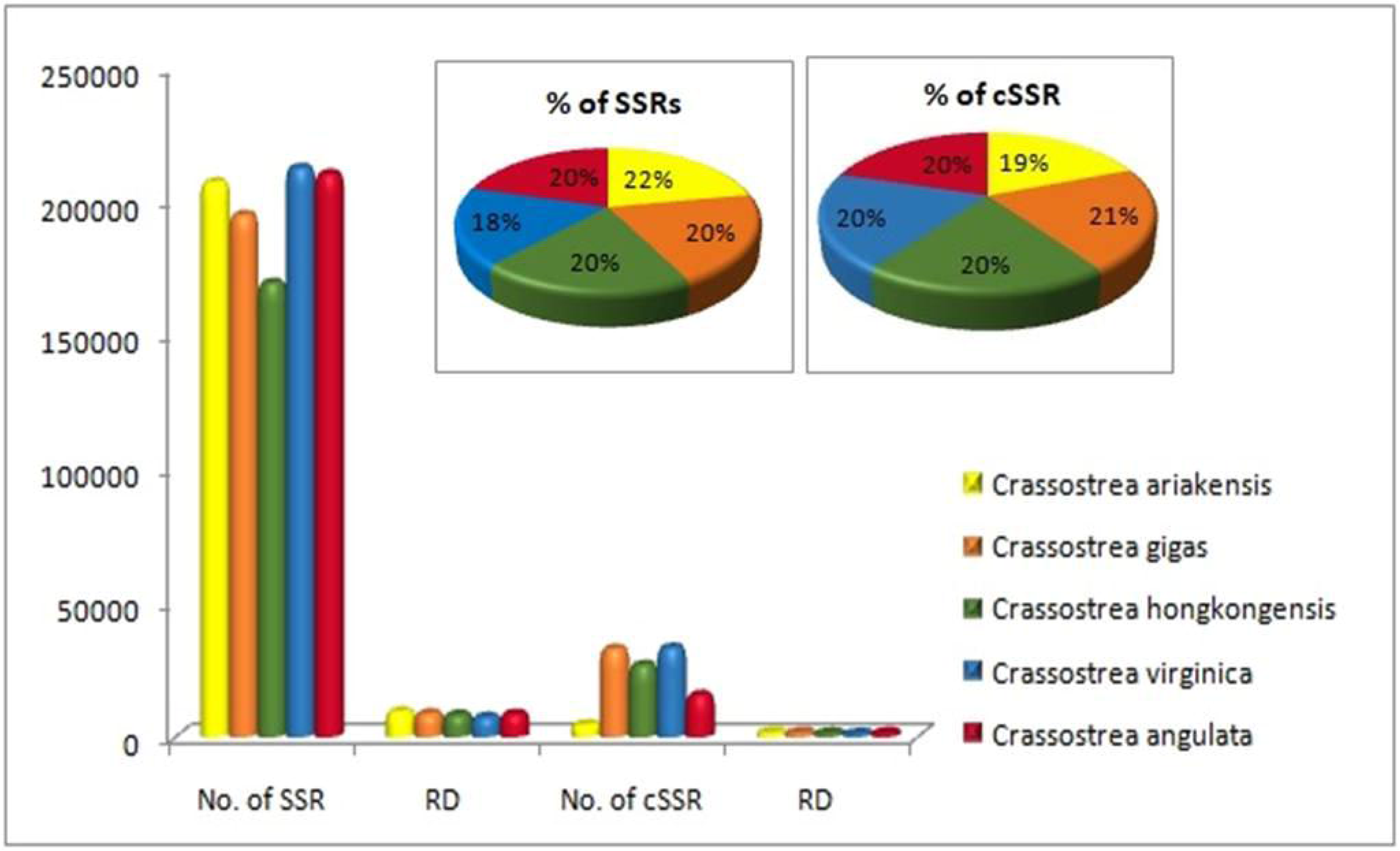
Graphical representation showing the distribution of SSR and cSSR frequency, RD, and percentage varied in the whole genome of different Crassostrea species.

**Figure 2:**
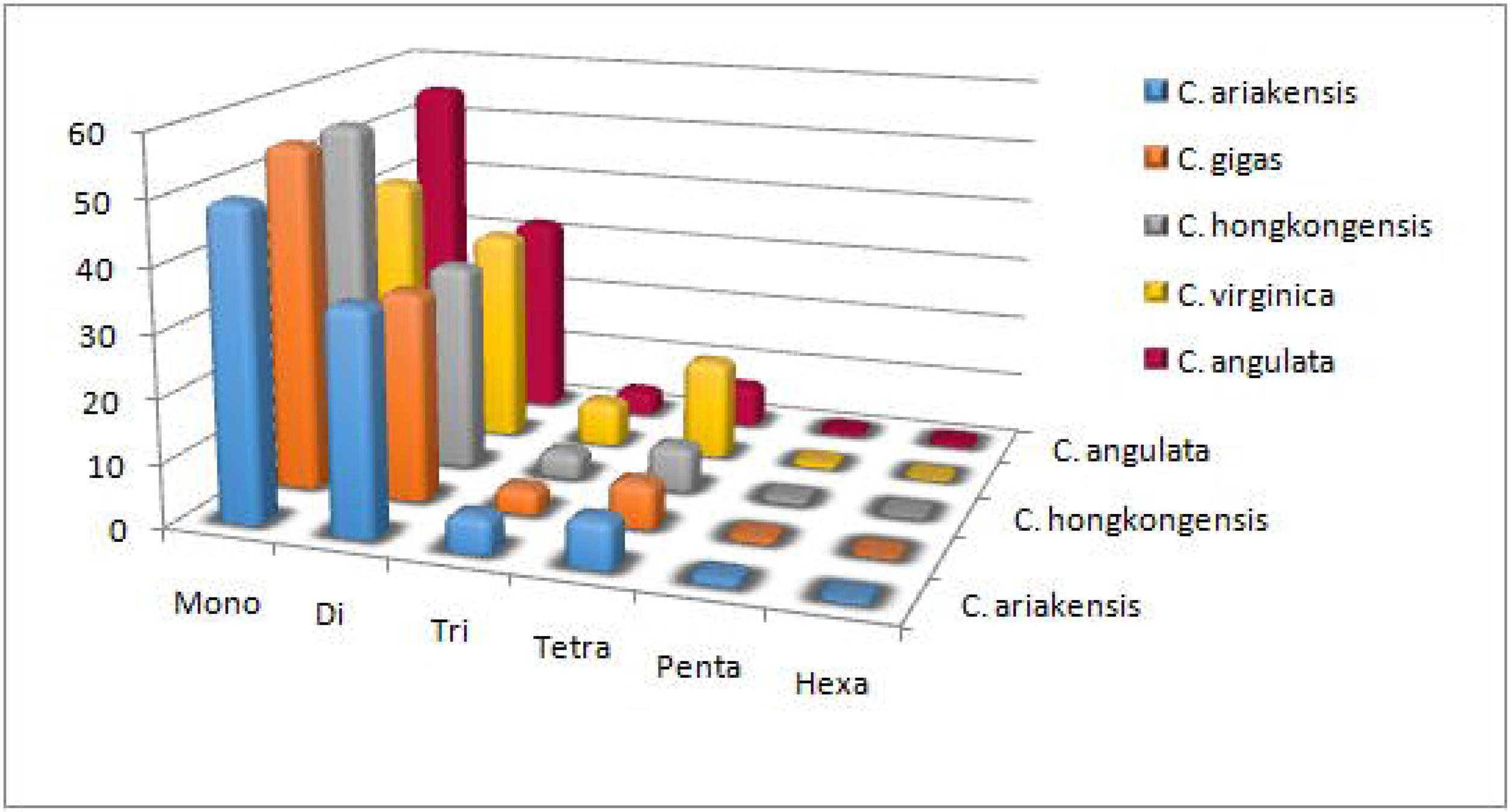
SSR percentage of different motifs (mono-hexa) in the complete genome.

**Figure 3:**
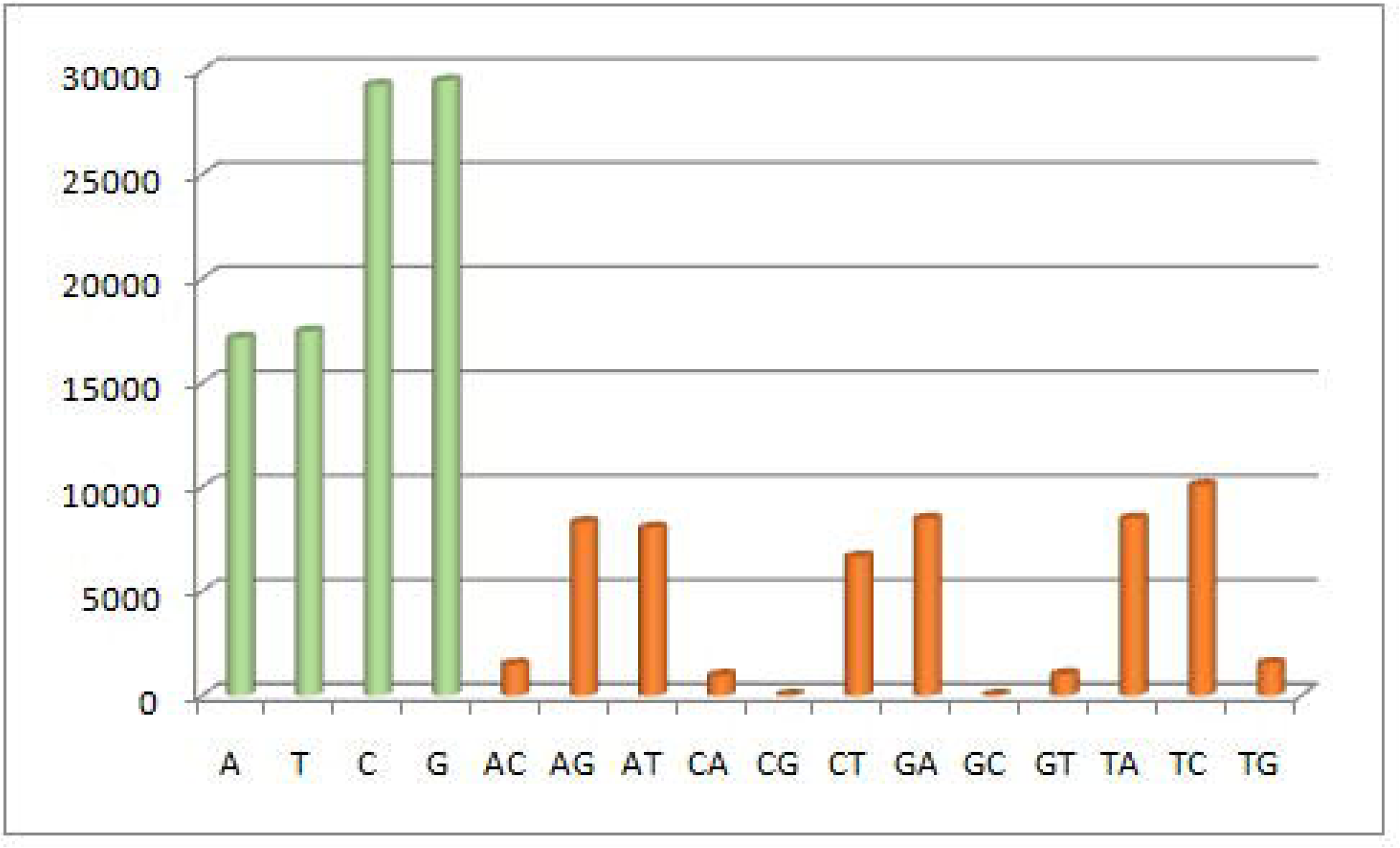
Overall SSR repeat motifs in *C. hongkongensis* throughout the genome. **A.** Mono and dinucleotide repeat motifs showing C/G, AG, AT, TA, TC richness in the genome. **B.** Trinucleotide repeat motif showing AAT, TGT richness across the genome.

### 3.2. Chromosomes-wide distribution of perfect microsatellites

The chromosomes-wide distribution of five Crassostrea species revealed that *C. virginica* has the highest number of SSR in the majority of chromosomes (chr. 5, 8, and 9) followed by *C. ariakensis* (chr. 1, 3 and 4) and *C. angulata* (chr. 2, 7, and 10). The lowest number of microsatellites were observed in chromosome 5 with 29050 loci and 25850 loci in *C. virginica* and *C. gigas* respectively, followed by chromosome 2, 8, 3, and 1. The lowest frequency of microsatellites was found in chromosome 10 with 9376 and 11565 loci of *C. virginica* and *C. hongkongensis*, respectively (Table-2, Fig. 4). Highest RA was seen in *C. angulata* (206.2) in chromosome 9 followed by 204.31 (*C. hongkongensis*), 203.41 (*C.angulata*) in chromosome 10 and 2, respectively (Fig. 5). Similarly the RD was significantly correlated with RA values where maximum RA has more RD except chromosome 8, 9, and 10. The maximum RD was 11029.56 in chromosome 9 followed by 10241.84 and 10034.02 of *C. gigas* in chromosomes 8 and 2 respectively (Fig. 6).

**Figure 4:**
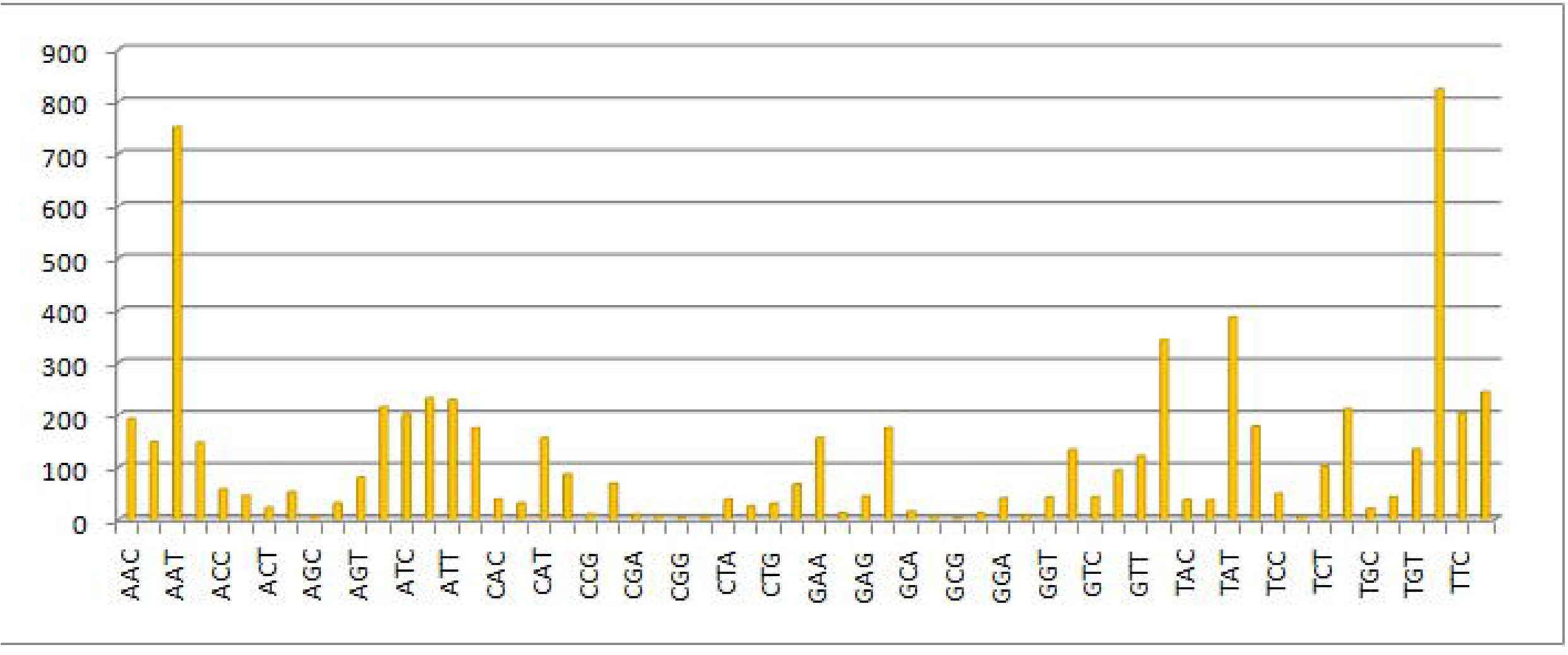
Characterization and distribution of SSR frequency across entire Chromosomes in all species.

**Figure 5:**
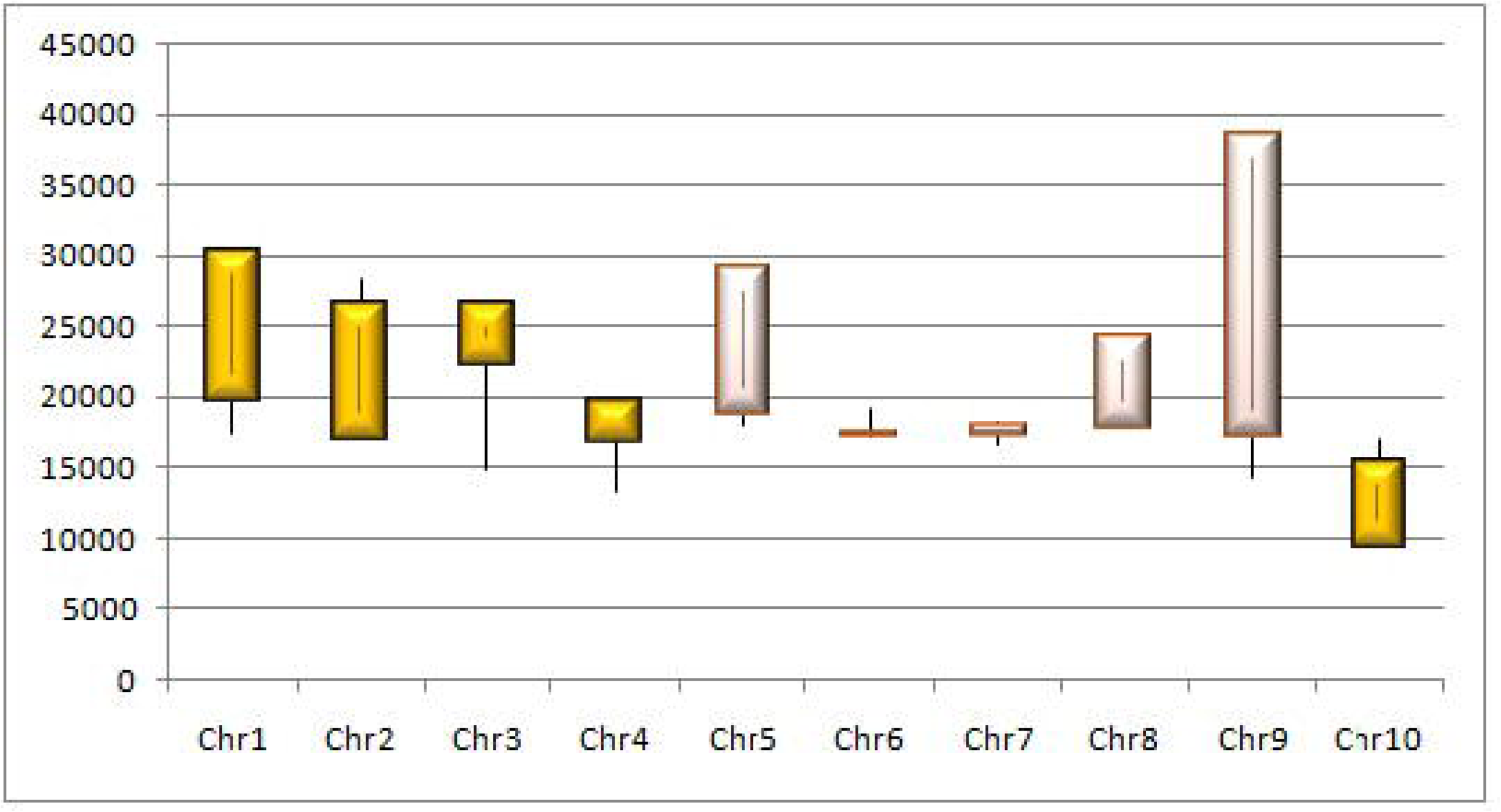
Comparison of relative abundance of SSRs from monomer to hexamer in all 5 Crassostrea chromosomes set.

**Figure 6:**
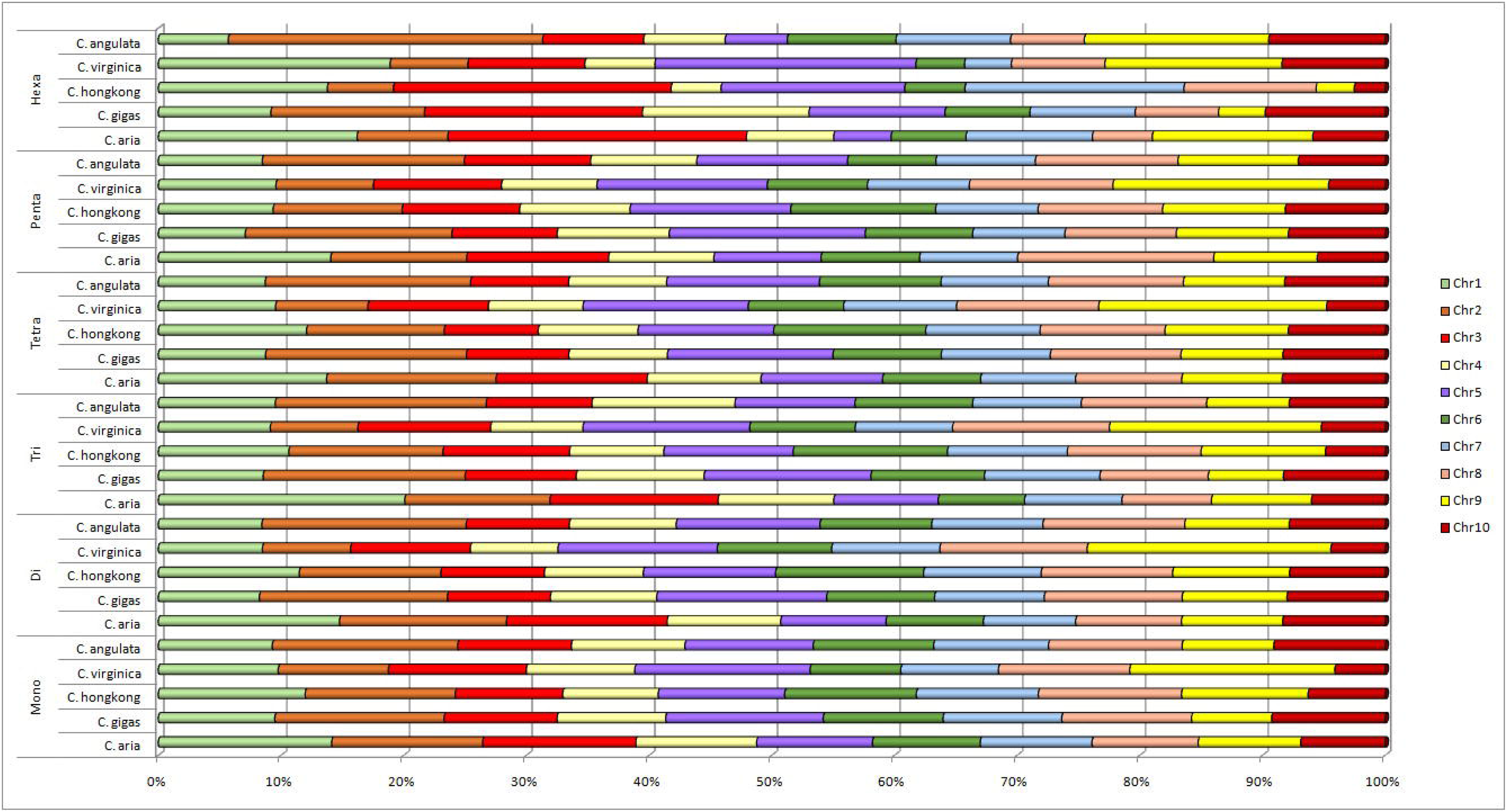
Relative density of mono-hexamer repeats across the entire Crassostrea species.

**Table 2:** The enrichment of perfect SSR and cSSR in chromosomes along with relative abundance, relative density, and percentage among five Crassostrea species.

|  | Chr. 1 | Chr. 2 | Chr. 3 | Chr. 4 | Chr. 5 | Chr. 6 | Chr. 7 | Chr. 8 | Chr. 9 | Chr. 10 |
| --- | --- | --- | --- | --- | --- | --- | --- | --- | --- | --- |
| *C. angulata* |  |  |  |  |  |  |  |  |  |  |
| SSR | 17725 | 31335 | 17557 | 17940 | 21703 | 18886 | 18176 | 21915 | 15572 | 16895 |
| RA | 307.7 | 383.89 | 301.95 | 346.2 | 350.4 | 305.8 | 298.7 | 362.4 | 391.14 | 300.5 |
| RD | 7781.31 | 9751.71 | 7513.35 | 8874.0 | 9510.3 | 7796.9 | 7550.71 | 9576.9 | 10623.8 | 7389.3 |
| % SSR | 0.78 | 0.98 | 0.76 | 0.89 | 0.96 | 0.78 | 0.76 | 0.96 | 1.07 | 0.74 |
| cSSR | 1264 | 2548 | 1193 | 1305 | 1795 | 1343 | 1228 | 1688 | 1287 | 1089 |
| cRA | 21.94 | 31.22 | 20.52 | 25.19 | 28.99 | 21.75 | 20.18 | 27.92 | 32.33 | 19.37 |
| cRD | 1375.28 | 1933.34 | 1209.2 | 1597.7 | 1925.7 | 1384.4 | 1283.44 | 1786.0 | 2092.8 | 1144.05 |
| cSSR% | 15.4 | 17.54 | 14.36 | 15.8 | 17.9 | 15.15 | 14.28 | 16.42 | 18.01 | 13.65 |
| ***C. ariakensis*** |  |  |  |  |  |  |  |  |  |  |
| **SSR** | 30365 | 26485 | 26543 | 19735 | 18776 | 17167 | 17310 | 17867 | 17233 | 15406 |
| **RA** | 352.46 | 329.83 | 334.11 | 282.08 | 283.09 | 277.94 | 285.25 | 299.11 | 310.08 | 368.83 |
| **RD** | 9491.05 | 9117.01 | 9142.25 | 7212.83 | 7152.59 | 6495.07 | 7014.5 | 7657.59 | 8118.51 | 10365.58 |
| **% SSR** | 0.95 | 0.92 | 0.92 | 0.73 | 0.72 | 0.7 | 0.71 | 0.77 | 0.82 | 1.04 |
| **cSSR** | 2145 | 2063 | 2044 | 1283 | 1161 | 1075 | 1062 | 1215 | 1206 | 1227 |
| **cRA** | 24.9 | 25.69 | 25.73 | 18.34 | 17.5 | 17.4 | 17.5 | 20.34 | 21.7 | 29.38 |
| **cRD** | 1754.6 | 1787.08 | 1818.93 | 1224.82 | 1137.21 | 1137.52 | 1134.15 | 1394.53 | 1523.46 | 1993.99 |
| **cSSR%** | 15.31 | 16.82 | 16.6 | 13.93 | 13.11 | 13.4 | 13.05 | 14.63 | 15.14 | 17.15 |
| ***C. gigas*** |  |  |  |  |  |  |  |  |  |  |
| **SSR** | 17464 | 28455 | 17344 | 17163 | 25850 | 18198 | 18195 | 20852 | 14251 | 16996 |
| **RA** | 313.07 | 388.63 | 297.41 | 323.06 | 351.47 | 302.55 | 292.97 | 356.68 | 384.84 | 295.37 |
| **RD** | 814.63 | 10034.02 | 7664.65 | 8560.6 | 9745.01 | 7811.14 | 7524.88 | 10241.84 | 11029.59 | 7473.27 |
| **% SSR** | 0.82 | 1.01 | 0.77 | 0.86 | 0.98 | 0.79 | 0.76 | 1.03 | 1.11 | 0.75 |
| **cSSR** | 1307 | 2476 | 1244 | 1292 | 2094 | 1341 | 1265 | 1671 | 1202 | 1238 |
| **cRA** | 23.43 | 33.82 | 21.33 | 24.87 | 28.47 | 22.29 | 20.37 | 28.58 | 32.41 | 21.52 |
| **cRD** | 1662.94 | 2030.76 | 1354.83 | 1581.92 | 1933.82 | 1430.7 | 1277 | 2384.1 | 2236.93 | 1343.23 |
| **cSSR%** | 17.59 | 18.85 | 15.5 | 16.27 | 17.55 | 15.78 | 14.93 | 17.57 | 18.44 | 15.42 |
| ***C. hongkongensis*** |  |  |  |  |  |  |  |  |  |  |
| **SSR** | 19915 | 20158 | 14773 | 13370 | 17901 | 19208 | 16609 | 18990 | 16943 | 11565 |
| **RA** | 308.91 | 315.61 | 329.87 | 372.32 | 346.89 | 345.31 | 297.36 | 302 | 295.2 | 402.86 |
| **RD** | 7933.2 | 7881.63 | 8487.76 | 9321.07 | 9165.83 | 9351.43 | 7438.27 | 7452.47 | 7326.24 | 10969.85 |
| **% SSR** | 0.8 | 0.79 | 0.85 | 0.94 | 0.92 | 0.94 | 0.75 | 0.75 | 0.74 | 1.1 |
| **cSSR** | 1491 | 1450 | 1064 | 1004 | 1380 | 1570 | 1187 | 1278 | 1126 | 958 |
| **cRA** | 23.13 | 22.7 | 23.76 | 27.96 | 26.74 | 28.22 | 21.25 | 20.32 | 19.62 | 33.37 |
| **cRD** | 1512.88 | 1368.9 | 1633.95 | 1687.51 | 1806.95 | 1927.73 | 1341.39 | 1263.7 | 1206.18 | 2134.63 |
| **cSSR%** | 16.03 | 15.22 | 15.9 | 16.06 | 16.69 | 17.66 | 15.54 | 14.38 | 14.13 | 17.96 |
| ***C. virginica*** |  |  |  |  |  |  |  |  |  |  |
| **SSR** | 19710 | 16959 | 22242 | 16937 | 29050 | 17400 | 17924 | 24276 | 38494 | 9376 |
| **RA** | 300.17 | 274.65 | 288.65 | 283.77 | 294.35 | 339.51 | 309.97 | 319.69 | 369.57 | 287.21 |
| **RD** | 7623.27 | 6486.55 | 6869.73 | 6625.76 | 7068.25 | 8453.86 | 7705.98 | 7889.14 | 9326.1 | 6810.28 |
| **% SSR** | 0.73 | 0.65 | 0.69 | 0.67 | 0.71 | 0.85 | 0.78 | 0.79 | 0.94 | 0.69 |
| **cSSR** | 1308 | 4054 | 1450 | 1028 | 1956 | 1357 | 1358 | 1762 | 3238 | 699 |
| **cRA** | 19.92 | 17.07 | 18.82 | 17.22 | 19.82 | 26.48 | 23.48 | 23.2 | 31.09 | 21.41 |
| **cRD** | 1193.88 | 961.83 | 1064.11 | 968.51 | 1181.36 | 1537.22 | 1449.38 | 1449.61 | 1957.88 | 1342.41 |
| **cSSR%** | 14.3 | 13.28 | 14.14 | 13.1 | 14.74 | 16.69 | 16.71 | 16.21 | 18.84 | 16.69 |

#### 3.2.1. Mononucleotide repeats

The most predominant mononucleotide repeat motif is (C)n with an average value of 5143.7-6803.3 in chromosomes of five oyster species. The highest (C)n repeat was seen in *C. gigas* (6803.3) accounting for 63.9% of the total mononucleotide of all the chromosomes followed by *C. angulata* (6747.4), *C. virginica* (6229.8), *C. hongkongensis* (5877.7) and least in *C. ariakensis* (5143.7) which is 62.9% of the total mononucleotide. In contrary, (A)n was the less frequent repeat motif with the average of 2660.8, 3462.5, 3839.5, 3913.3, and 5070.4 occupying 29.9%, 36.07%, 36.7%, 37.1% and 49.6% of total mononucleotide SSRs in *C. virginica*, *C. gigas*, *C. angulata*, *C. hongkongensis*, and *C. ariakensis,* respectively. The most mononucleotide repeat times are 5437 (chr. 8), 4347 (chr. 6), 448 (chr. 2), 96 (chr. 4), and 75 (chr. 7) in *C. gigas*, *C. hongkongensis*, *C. virginica*, *C. virginica*, and *C. ariakensis*.

#### 3.2.2. Dinucleotide repeats

(AG)n repeat motif with the frequency of 3356.1-5863.1 loci/Mb was the most prevalent dinucleotide SSR occupying over 81.1%, 62.14%, 61.60%, 60.85%, and 57.40% of the total dinucleotide in *C. virginica*, *C. angulata*, *C. gigas*, *C. hongkongensis* and *C. ariakensis* chromosomes, respectively. The second most predominant repeat motif is (AT)n with a frequency of 873.7-2430.3 loci/Mb accounting 33.3, 30, 29.9, 29.32, and 12.08% in *C. ariakensis*, *C. gigas*, *C. hongkongensis*, *C. angulata*, and *C. virginica,* respectively. *C. virginica* had a relatively low frequency of (AT)n compared to other species. (AT)n found to be most predominant on all Crassostrea species with the highest 3503 in *C. virginica* (chr. 9), 2039 in *C. ariakensis* (chr. 1), and 2395, 2154, 1174 in *C. gigas*, *C. angulata*, *C. hongkongensis* within chromosome 2 of each animals. The second most frequent dinucleotide SSR was (TA)n in four species except *C. virginica* with the mean 1501.5 (Fig. 7). The utmost dinucleotide repeat times were 616 (chr. 1), 473 (chr. 5), 430 (chr. 8), 132 (chr. 6) and 115 (chr. 8) within *C. angulata*, *C. gigas, C. hongkongensis*, *C. virginica*, and *C. ariakensis,* respectively.

**Figure 7:**
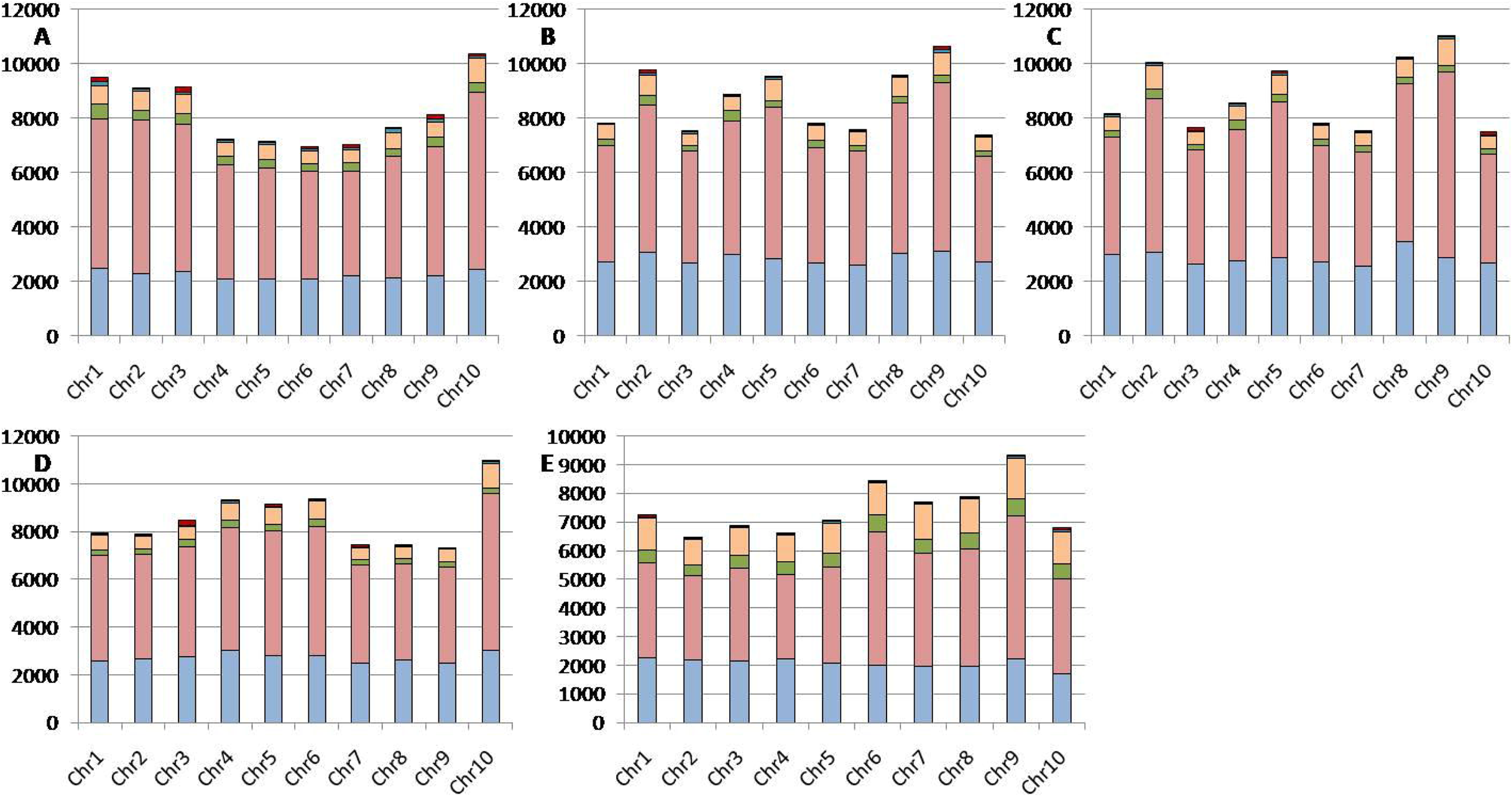
Loci/Mb heatmap for dinucleotide repeat in all Crassostrea species such as (A) *C. ariakensis*, (B) *C. angulata*, (C) *C. gigas*, (D) *C. hongkongensis*, (E) *C. virginica*. Colors from red to blue indicated the highest to lowest number of dinucleotide repeat motifs.

#### 3.2.3. Trinucleotide repeats

A total of 48918 trinucleotide repeats were found in all five Crassostrea chromosomes with the highest frequency in *C. virginica* (14998) and the lowest in *C. hongkongensis* (6822). The most abundant trinucleotide repeat (AAT)n that accounts for 56.67%, 40.20%, 39.51%, 39.36%, and 28.71% in *C. ariakensis*, *C. hongkongensis*, *C. angulata*, *C. gigas,* and *C. virginica,* respectively, followed by (ATC)n, (AAC)n, (AAG)n. The least frequent trinucleotide repeat motifs were (AGC)n and (CCG)n with a frequency of less than 0.4 loci/Mb. The highest 1684 loci/Mb (AAT)n trinucleotide repeat was found in chromosome 1 of *C. ariakensis* which occupied 14.3% of total trinucleotide repeat (Fig. 8). The highest trinucleotide repeat times were 53 (chr. 8), 49 (chr. 4), 47 (chr.10), 44 (chr. 1), and 43 (chr. 2) in *C. hongkongensis*, *C. angulata*, *C. gigas*, *C. virginica*, and *C. ariakensis,* respectively.

**Figure 8:**
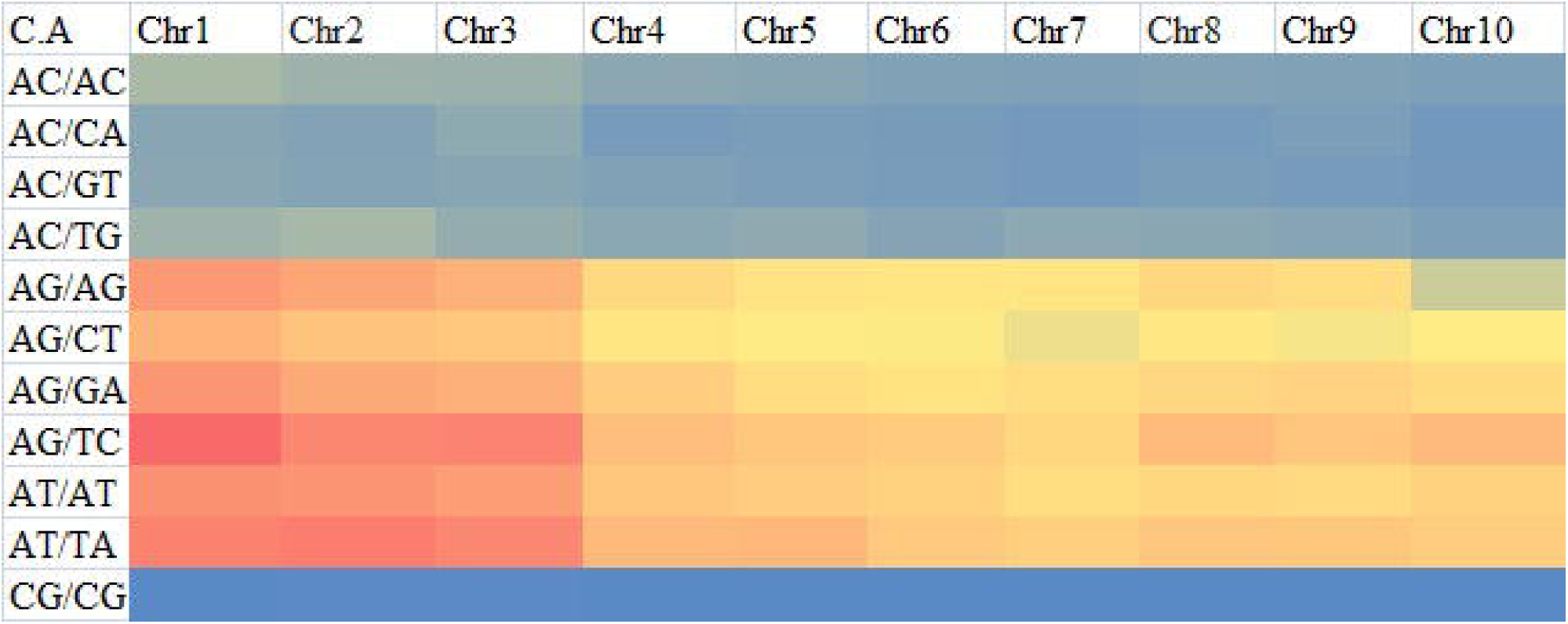
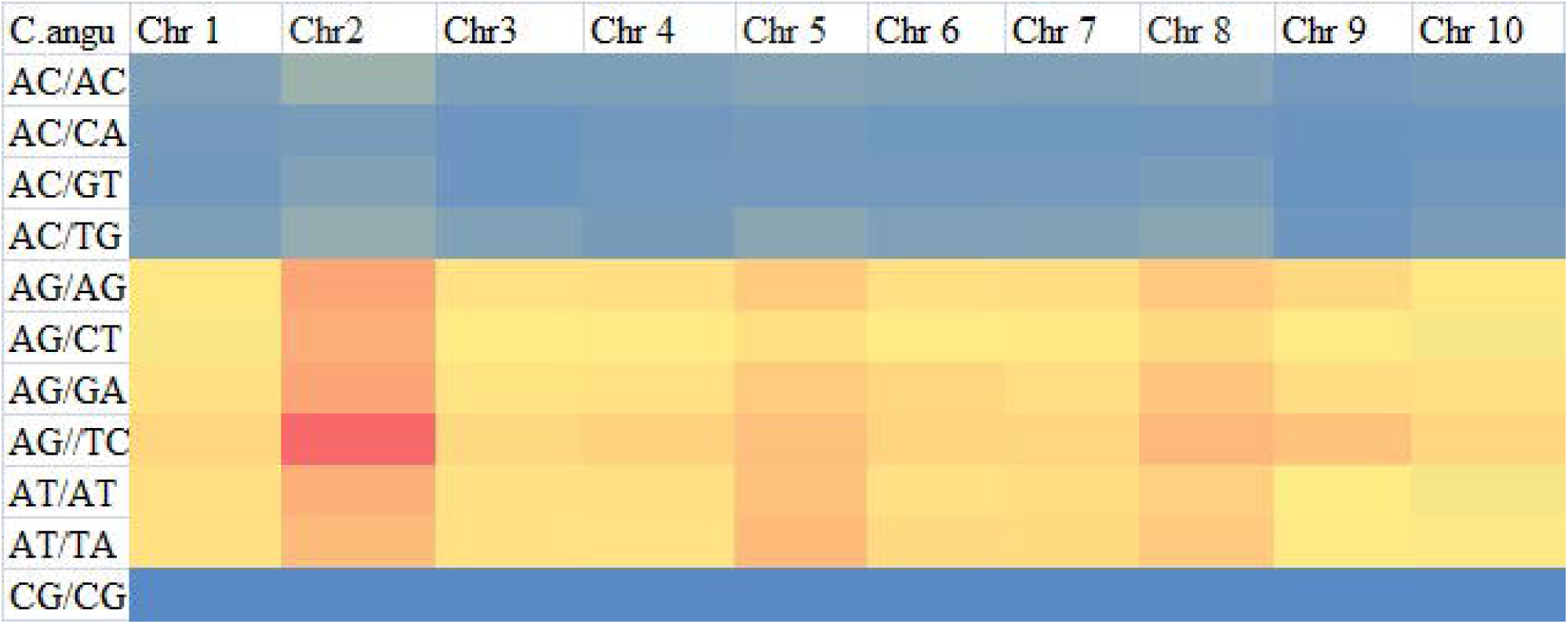
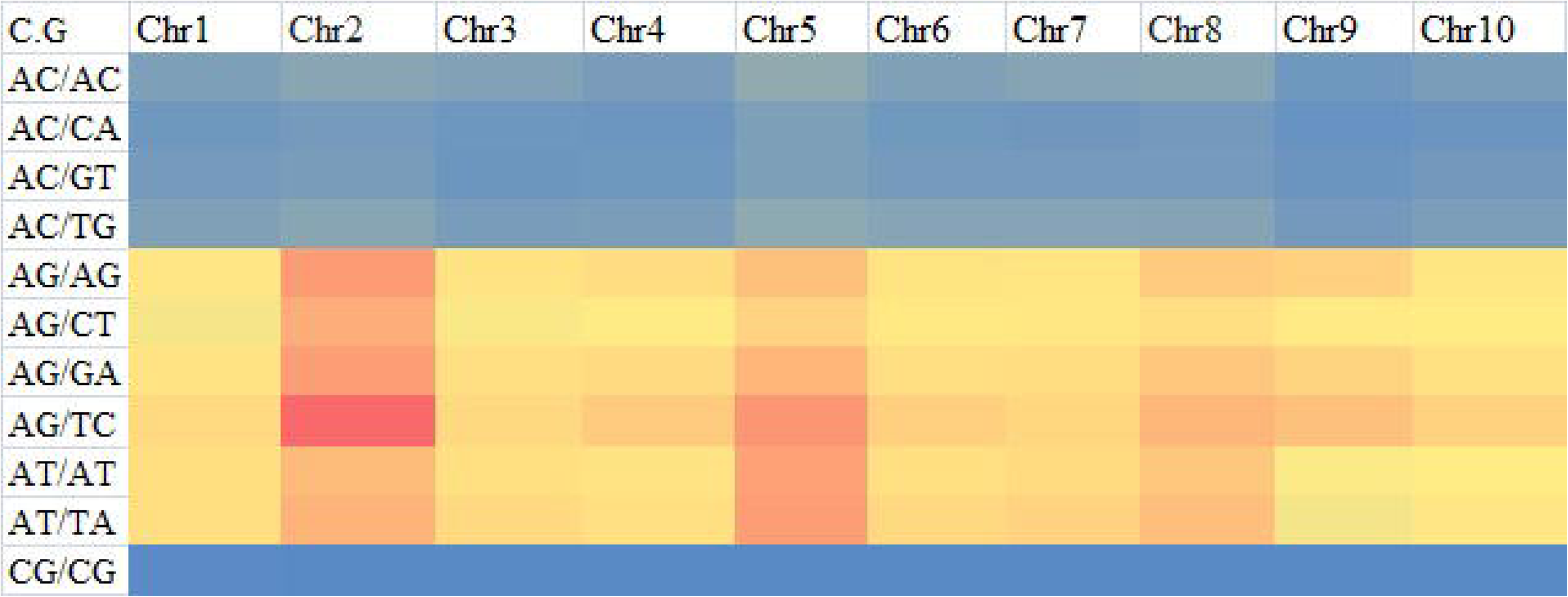
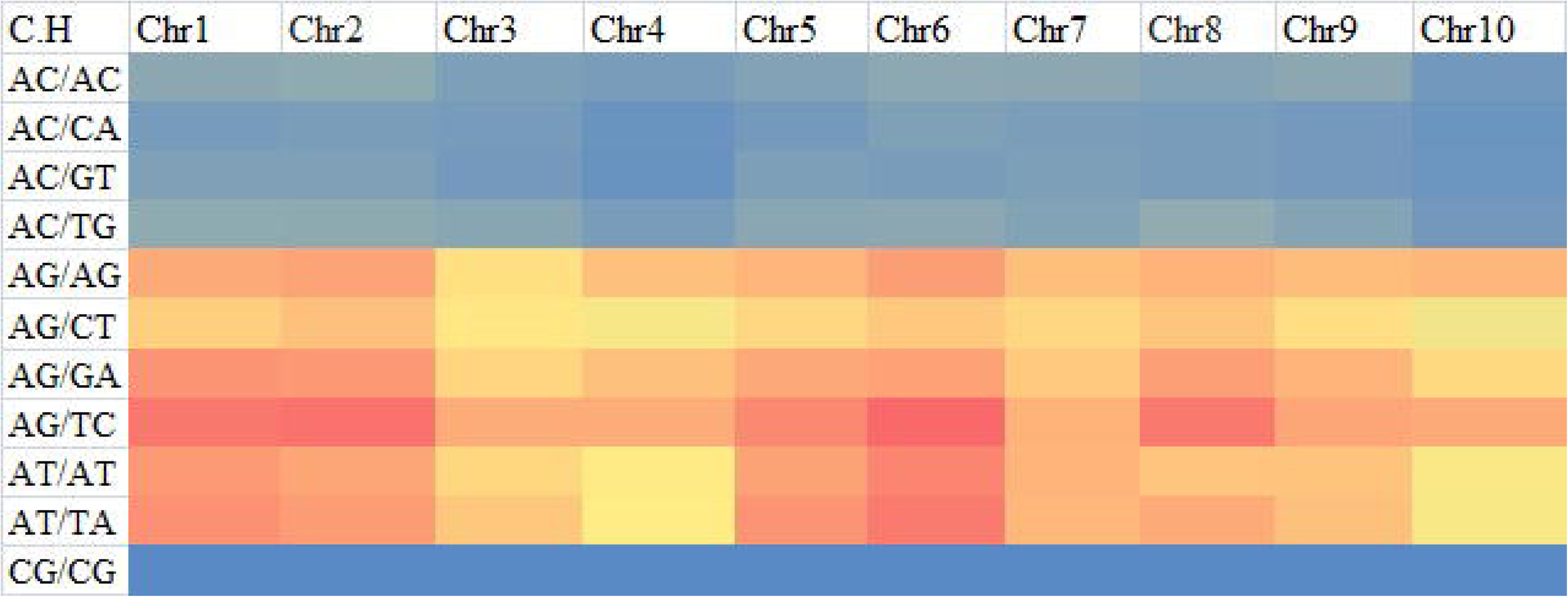
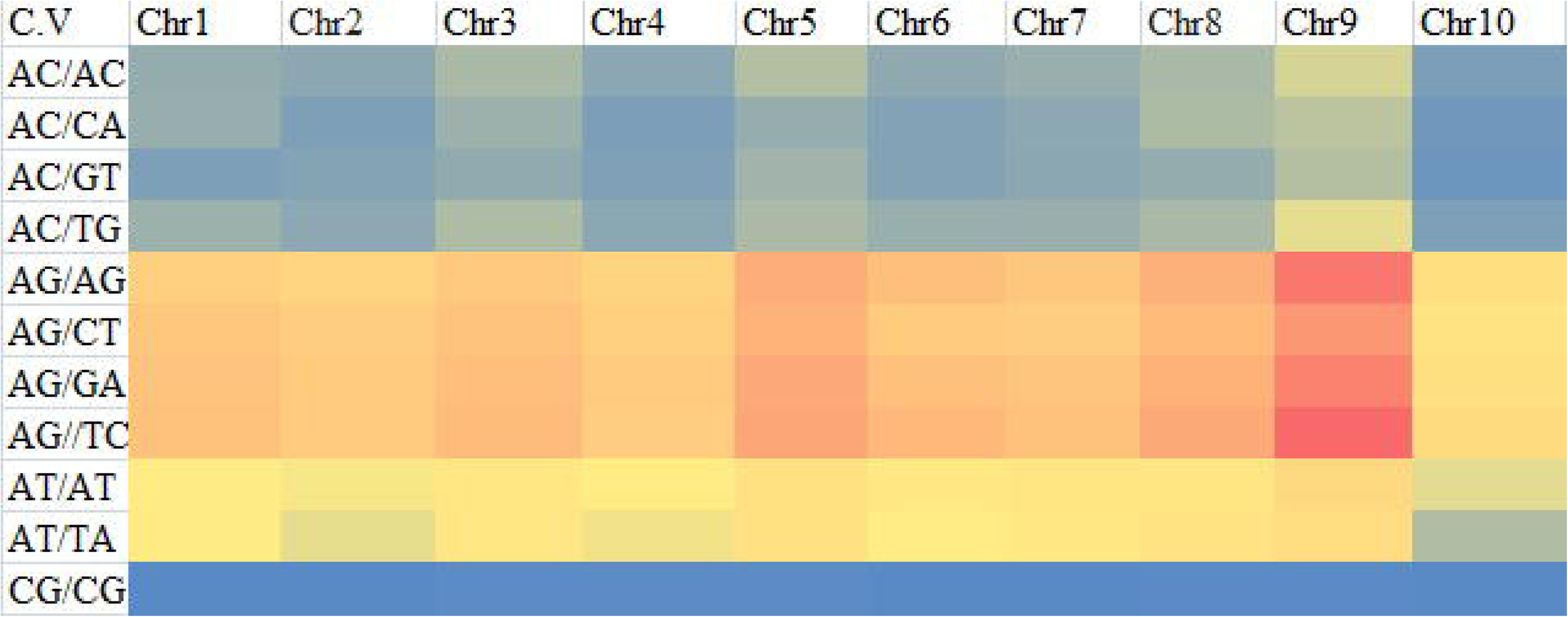
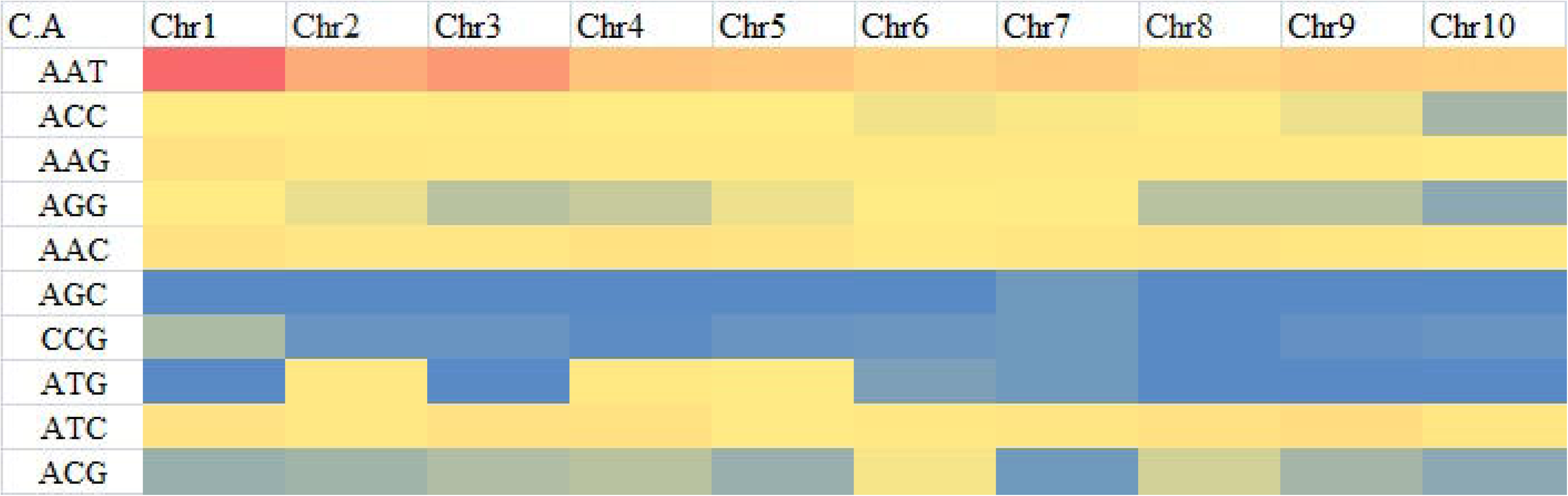
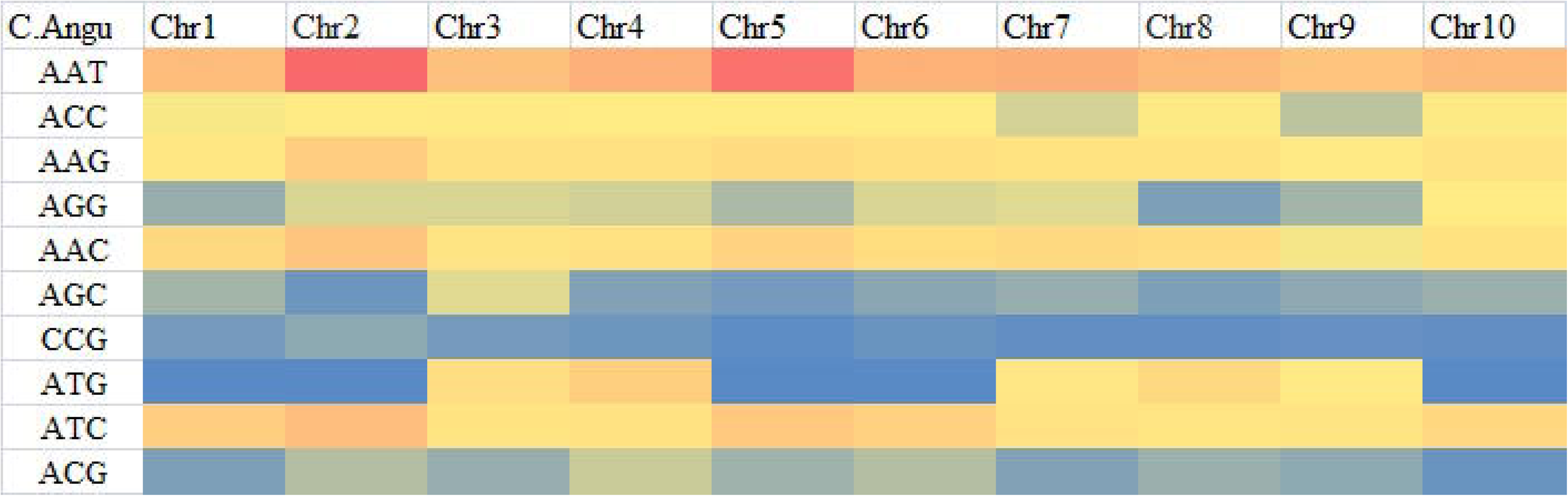
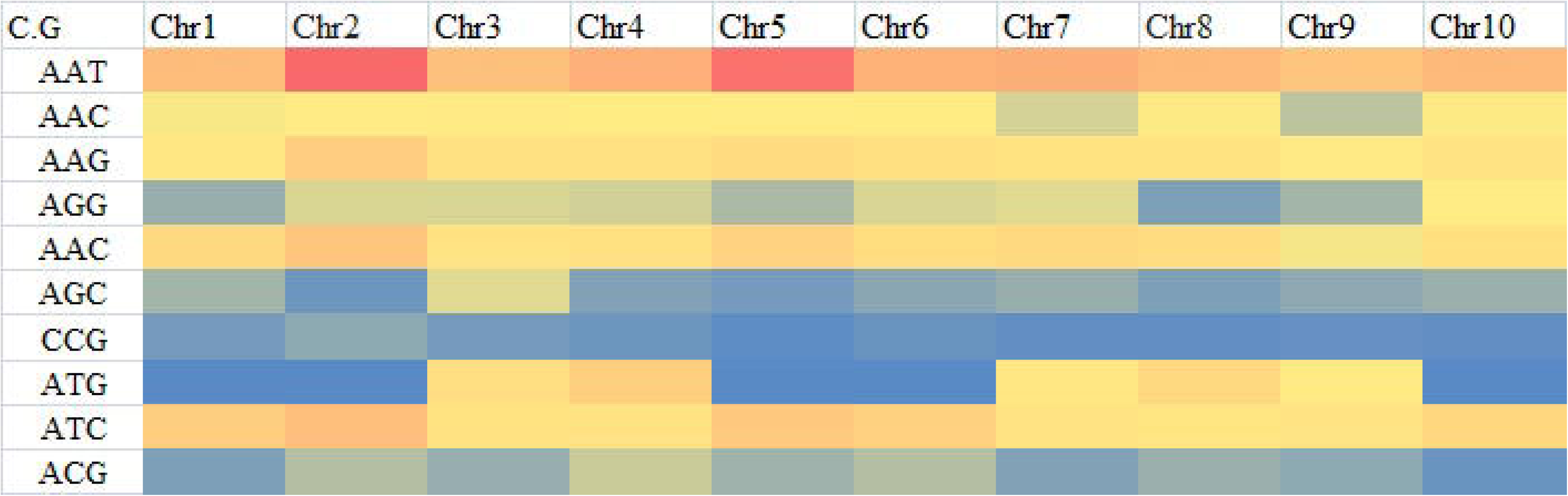
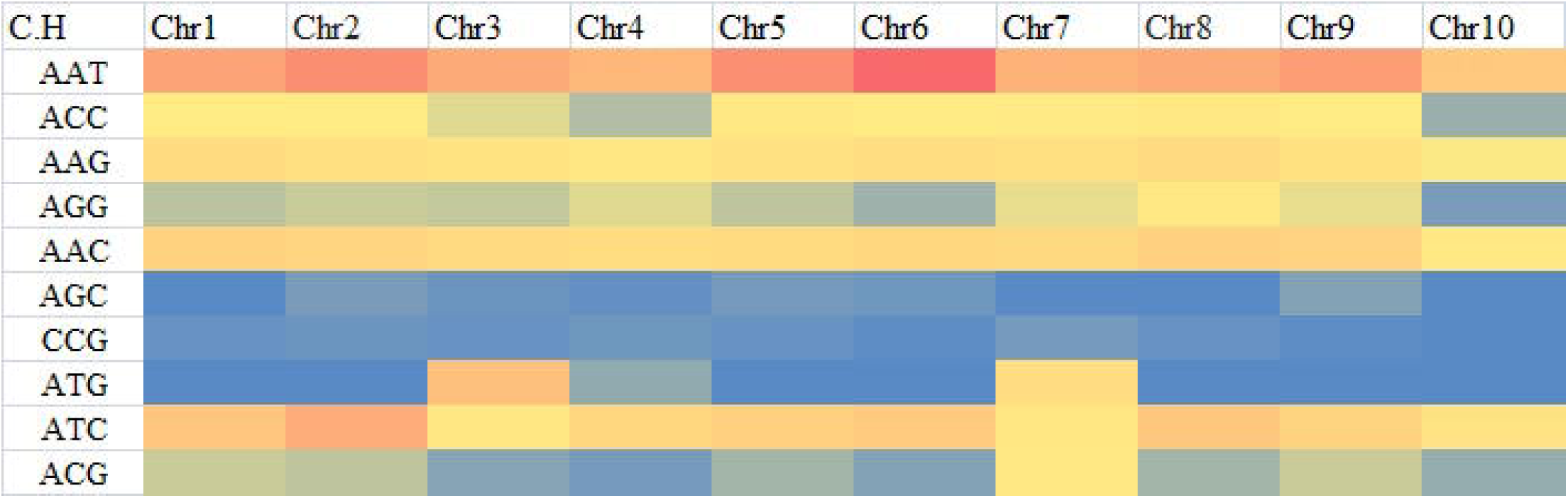
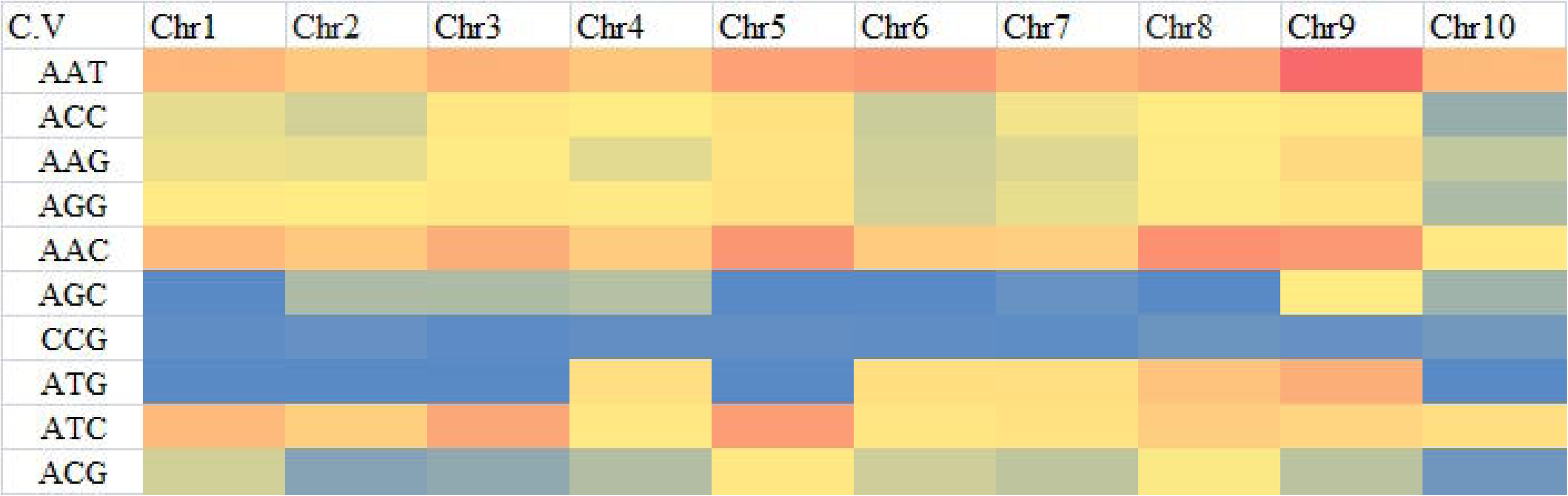
Heatmap of trinucleotide repeat motifs showed the relative numbers among chromosomes of all five Crassostrea species such as (A) *C. ariakensis*, (B) *C. angulata*, (C) *C. gigas*, (D) *C. hongkongensis*, (E) *C. virginica*. Color key from red to blue indicated numbers per Mb in descending order.

#### 3.2.4. Tetranucleotide repeats

(AAAC)n repeat with frequencies of 309.3 loci/Mb and 309.5 loci/Mb were the most frequent tetranucleotide microsatellites occupying 22.98 and 22.56 % of the total number of tetranucleotide microsatellites in *C. angulata* and *C. gigas*, respectively. Whereas (ACAG)n, (ACGG)n, and (AATC)n tetranucleotides were highly found in *C. ariakensis*, *C. hongkongensis*, *C. virginica,* respectively. The maximum of tetranucleotide repeat times was 219 (chr. 3), 184 (chr. 7), 174 (chr. 10), 166 (chr. 2), and 124 (chr. 9) in *C. ariakensis*, *C. hongkongensis*, *C. angulata*, *C. gigas* and *C. virginica,* respectively.

#### 3.2.5. Pentanucleotide repeats

The pentanucleotide motif (AAAAC)n was predominantly with average frequencies of 19.8–33 loci/Mb. The highest average frequency was observed in *C. ariakensis* (330) followed by *C. gigas* (257), *C. angulata* (249), *C. hongkongensis* (232) and least in *C. virginica* (198). The second most penta nucleotide repeat motif was (ATAGG)n with the highest average frequency of 34.4 loci/Mb in *C. gigas*. The highest observed pentanucleotide repeat times were 66 (chr. 7), 30 (chr. 5), 21 (chr. 3), 19 (chr. 3) and 18 (chr. 8) in *C. ariakensis*, *C. virginica*, *C. hongkongensis*, *C. gigas* and *C. angulata,* respectively.

#### 3.2.6. Hexanucleotide repeats

Hexanucleotide microsatellites had a lower frequency and density than other SSRs. The most predominant hexa-motif in all Crassostrea chromosomes was (AACCCT)n with the average frequency ranging from 21.6 (*C. gigas*)-72.8 (*C. ariakensis*) followed by (ACGCCG)n, (AAAAAC)n and (ACCCCC)n. (AACCCT)n repeat motif accounts for 54.64, 49.35, 42.53, 36.82, and 34.72% in *C. hongkongensis*, *C. ariakensis*, *C. virginica*, *C. gigas* and *C. angulata,* respectively. The most tetranucleotide repeat times were 299 (chr. 10), 122 (chr. 9), 81 (chr. 2), 51 (chr. 3), and 42 (chr.10) in *C. gigas*, *C. ariakensis*, *C. angulata*, *C. hongkongensis,* and *C. virginica,* respectively.

### 3.3. Regression analysis

The correlation and regression analysis revealed that the chromosome length of all the Crassostrea species had a positive and strong correlation with the incidence of SSR and cSSRs, but no significant influence on the GC content. In contrast, the number of cSSR had significant correlation with the GC content in *C. gigas* (*R^2^*=0.53; *P=*0.01), *C. hongkongensis* (*R^2^*=0.98; *P=*0.008) and *C. ariakensis* (*R^2^*=0.41; *P=*0.04) except *C. virginica* (*R^2^*=0.12; *P=*0.27) and *C. angulata* (*R^2^*=0.0001; *P=*0.09). Furthermore, we noticed that the chromosome length had no significant correlation with RA and RD as well as cRA and cRD in four oyster species, except *C. hongkongensis*. Similarly, GC content positively correlated with RA and RD as well as cRA and cRD in four species except *C. angulata*. The cSSR% was significantly correlated with the GC content in all five species, but no correlation was observed with chromosome size (Fig. 9).

**Figure 9:**
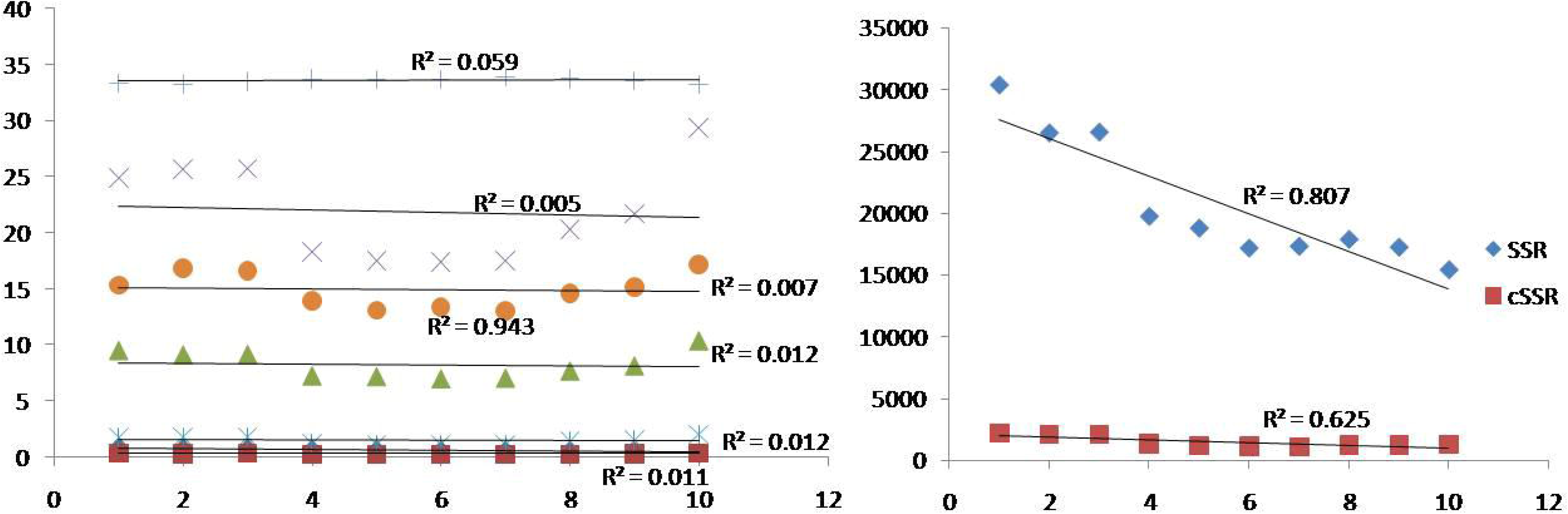
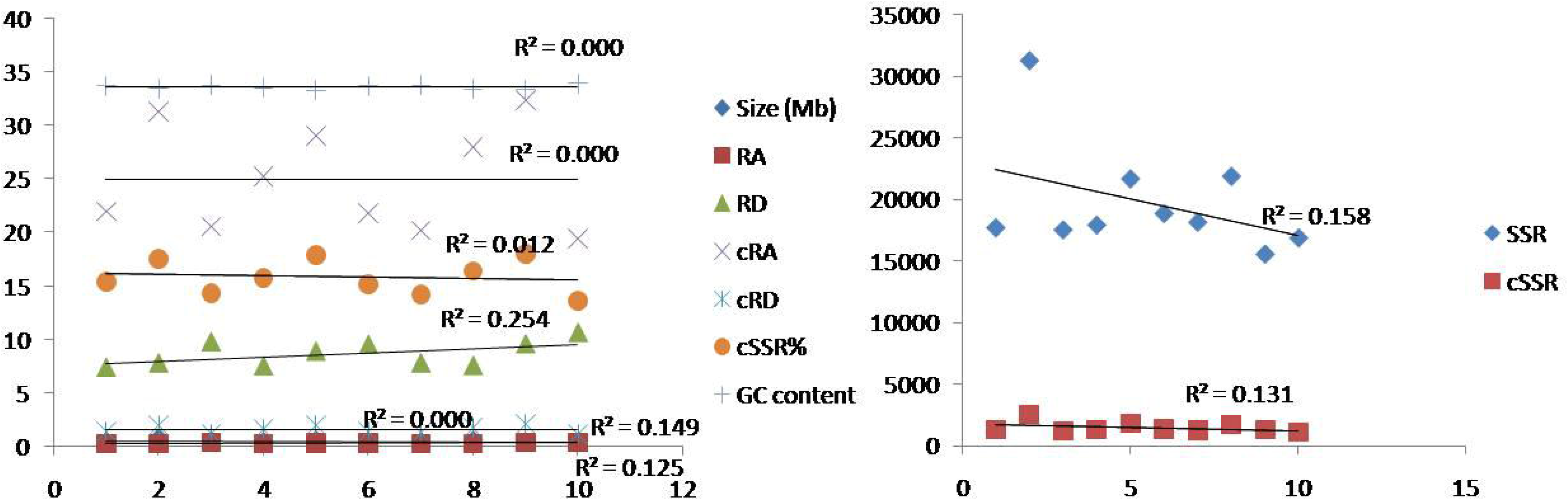
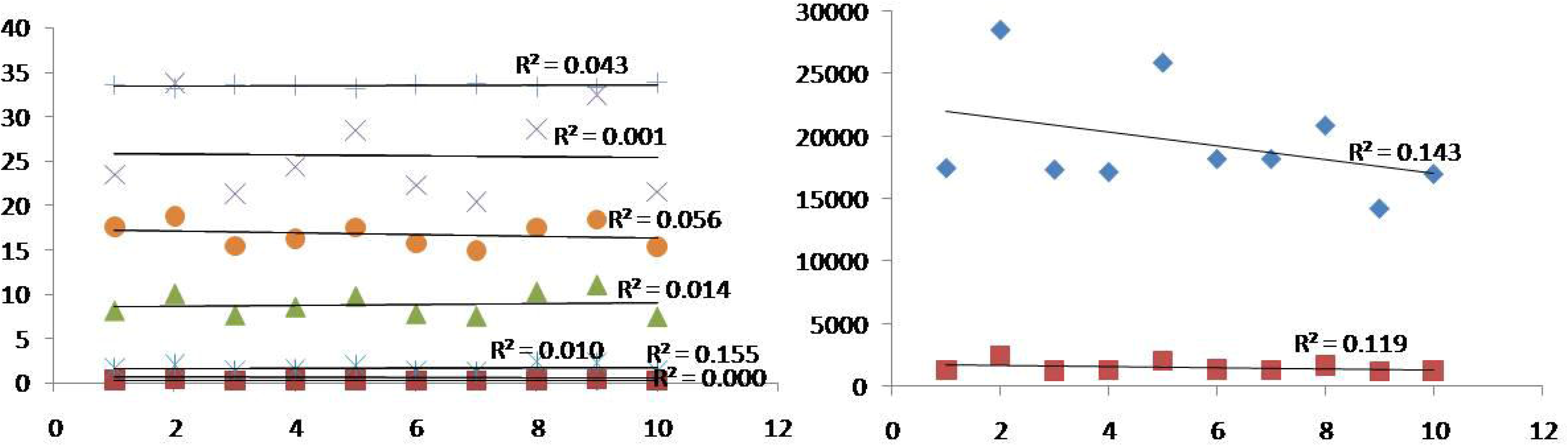
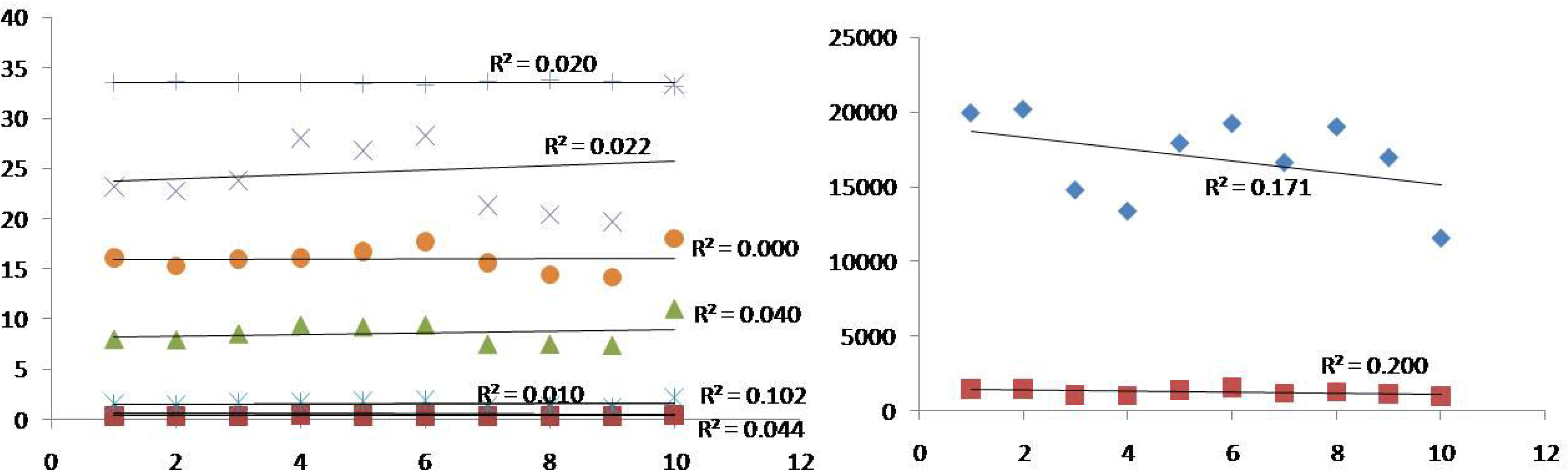
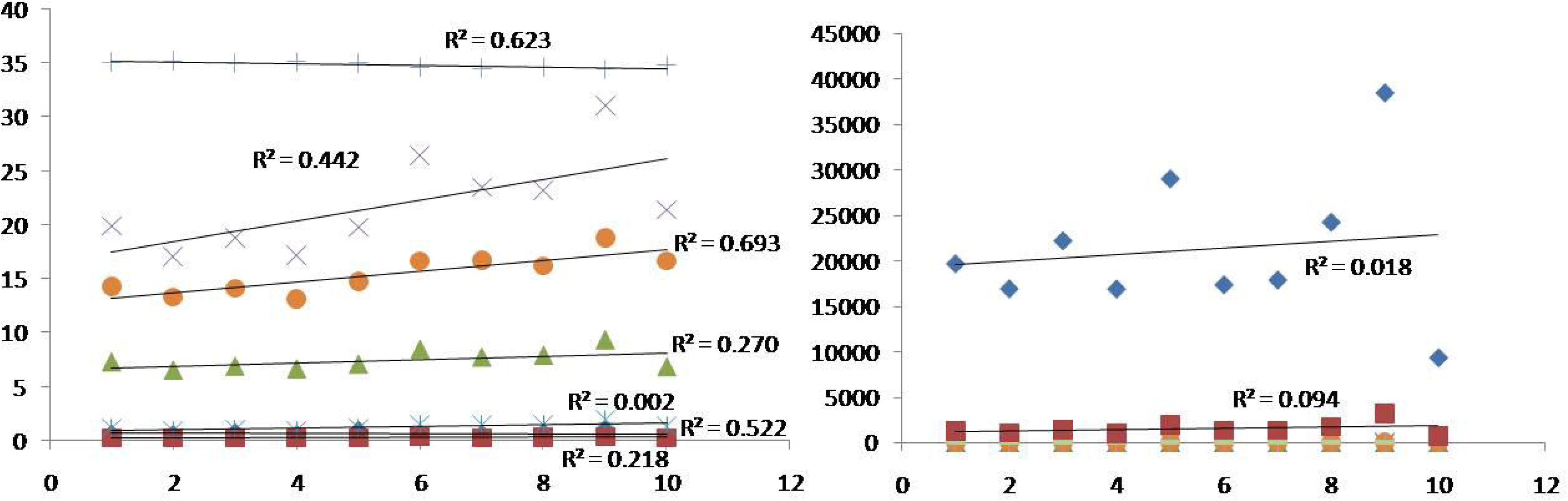
Association analysis between microsatellites and Crassostrea genomes. Correlation analysis of RA, RD, % SSR, cRA, cRD and % of cSSR with respect to genome size and GC content as well as regression analysis of SSR and cSSR with genome size and GC content of all five Crassostrea species. (A) *C. ariakensis*, (B) *C. angulata*, (C) *C. gigas*, (D) *C. hongkongensis*, (E) *C. virginica*. R^2^ represents the correlation coefficient and P< 0.05 was considered as the positive and significant correlation among them.

### 3.4. Occurrence of compound microsatellites

The genome-wide analysis of cSSR in five Crassostrea genomes resulted in a sum of 768733 microsatellites distributed throughout the genomes (Table 1). *C. ariakensis* (14436) had the lowest number of cSSR, whereas *C. gigas* (16520) had the highest. The cSSR% was found highest in *C. gigas* (16.75%), followed by *C. angulata* (16.23%), *C. hongkongensis* (16.08%), *C. virginia* (15.68%) and the lowest was in *C. ariakensis* (14.99%) (Fig. 1). The cRA ranged from 21.78-25.22 and noticed as highest in *C. angulata* and lowest in *C. ariakensis*. Similarly, cRD ranged from 1332.81–1694.54, with *C. gigas* having the highest followed by *C. angulata*, *C. hongkongensis*, *C. ariakensis,* and *C. virginica*. To understand if the cSSRs are distributed randomly throughout the genome or maintain regional bias, the clustering of cSSR was studied with the increasing of dMax from 10-100 using the Krait software. A significant upswing in cSSR% while increasing dMax suggested the random distribution of cSSR throughout the genome (Fig. 10).

**Figure 10:**
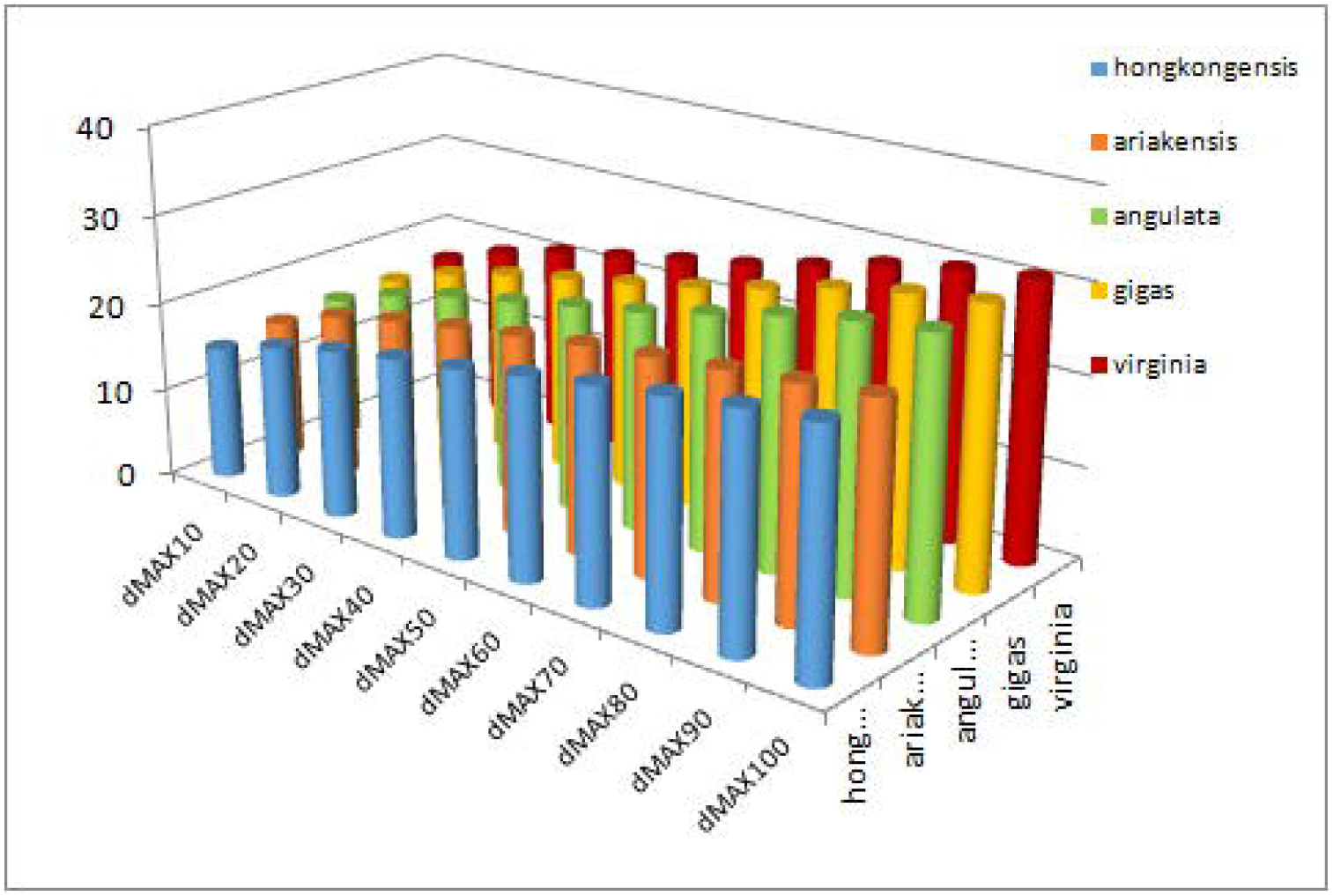
cSSR percentage with varying dMax (10-100) in the complete genome of all Crassostrea species.

The number of cSSR in the chromosomes of five Crassostrea species resulted highest in *C. gigas* in the majority of chromosomes (chr. 2, 5, and 6) followed by *C. angulata* (chr. 2, 5, and 8) and *C. virginia* (chr. 9, 5, and 8), *C. hongkongensis* (chr. 6, 1, 2) and *C. ariakensis* (chr. 1, 2, 3). The most frequent compound microsatellite were observed in chromosome 9 with 3238 loci and chromosome 2 with 2548 loci in *C. virginica* and *C. gigas* respectively, followed by chromosome 2 with 2476 loci in *C gigas*, chromosome 1 with 2145 loci in *C ariakensis* and chromosome 5 with 2094 in *C gigas*. The least frequent cSSR was found in chromosome 10 with 699 loci and 958 loci of *C. virginica* and *C. hongkongesnis,* respectively. The highest cSSR% was seen in *C. virginia* (18.85%) in chromosome 2 followed by Chromosome 2 (18.75%) and 9 (18.44%) of *C. gigas* respectively. The highest cRA was seen in *C. gigas* (33.82) in chromosome 2 followed by 33.37 (*C. hongkongensis*), 32.41 and 32.33 in chromosome 9 of *C. gigas* and *C. angulata,* respectively. The maximum cRD was 2384.1 and 2236.92 in chromosome 8 and 9 of *C. gigas* followed by 2134.63, 2092.87, and 2030.76 of *C. hongkongensis*, *C. angulata*, and *C. gigas* in chromosome 10, 9 and 2 respectively.

### 3.5. Motif complexity and structure of cSSR

Among all complete genomes, the most predominant cSSR motifs were (T)-X-(G) followed by (C)-X-(A), (TC)-X-(TC), (AG)-X-(AG), (GA)-X-(AG) and (C)-X-(AC). Other less frequent compound microsatellites were (T)-X-(T), (A)-X-(A), (C)-X-(C), (G)-X-(C), (G)-X-(G), (AC)-X-(AT), (AT)-X-(AT), (CT)-X-(CT), (GGGTTA)-X-(GGGGTTA), (TTTAGGG)-X-(GGGTTA), (GATA)-X-(GATA), (CT)-X-(CA) and (GA)-X-(CAGA) ubiquitously present in the genome. During motif duplication Similar motifs located on both ends with the spacer sequence is usually considered as duplicated cSSR such as (CA)n-(X)y-(CA)x. We observed a tremendous contribution of most frequently distributed duplicated cSSR, like (T)-X-(T), (A)-X-(A), (C)-X-(C), (G)-X-(C), (G)-X-(G), (AC)-X-(AT), (AT)-X-(AT), (CT)-X-(CT) towards genome evolution of the respective animals. The CTG–CAG cSSR composed of self-complementary motifs has been considered as a recombination hot spot (Liu, Chen et al. 2010). Our study did not observe such a motif suggesting that, CTG–CAG was not associated with recombination in Crassostrea. However, many cSSRs having other complementary motifs, such as (G)-X-(C), (AT)-X-(TA), and (CA)-X-(GT) were observed, which might be responsible for secondary structure formation.

### 3.6. Microsatellite characterization, diversity analysis, and cross-species amplification

The alleles number of the polymorphic cSSR loci developed during this study varied from 2-10, with an average of 4.44 alleles per locus (Table 3). Loci HKM-11 highest number of alleles (*Ne*=10) followed by HKM-19 (*Ne*=9) and HKM-9 (*Ne*=8). Observed heterozygosity (*Ho*) varied from 0.092-0.897, with an average of 0.631, while expected heterozygosity (*He*) ranged from 0.188-0.859, with a mean of 0.578 (Table 3). The exact tests for Hardy-Weinberg equilibrium (*HWE*), after Bonferroni correction deciphered that 22 of the 29 loci followed the equilibrium, and the rest seven loci deviated from HWE due to a decrease in heterozygosity. Additionally, seven pairs of loci (HKM-8, HKM-11-HKM-13, HKM-22, HKM-17) revealed significant linkage disequilibrium (*LD*) after Bonferroni correction. Polymorphic Information Content (*PIC*) of the 29 developed cSSR loci developed during this study ranged from 0.088-0.828, and the mean was 0.591 (Table 3). These results suggested that the population used during this study had good genetic diversity and the markers developed during this study were polymorphic enough to decipher genetic diversity. Furthermore inbreeding coefficient (*Fis*) was obtained that ranged from 0.0001-1 with an average value 0.416, which suggested the presence of moderate inbreeding within the analyses population. Cross-species amplification of the developed markers revealed that 15, 10, and 8 loci of amplified with *C. ariakensis* and *C. gigas*, and *C. angulata,* respectively (Table 4).

**Table 3:**
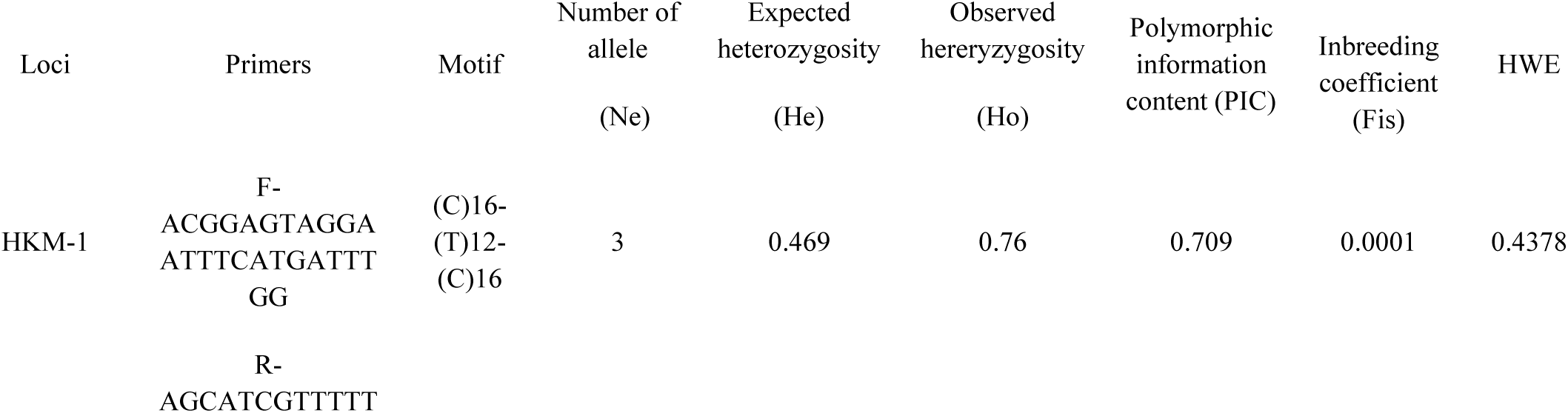
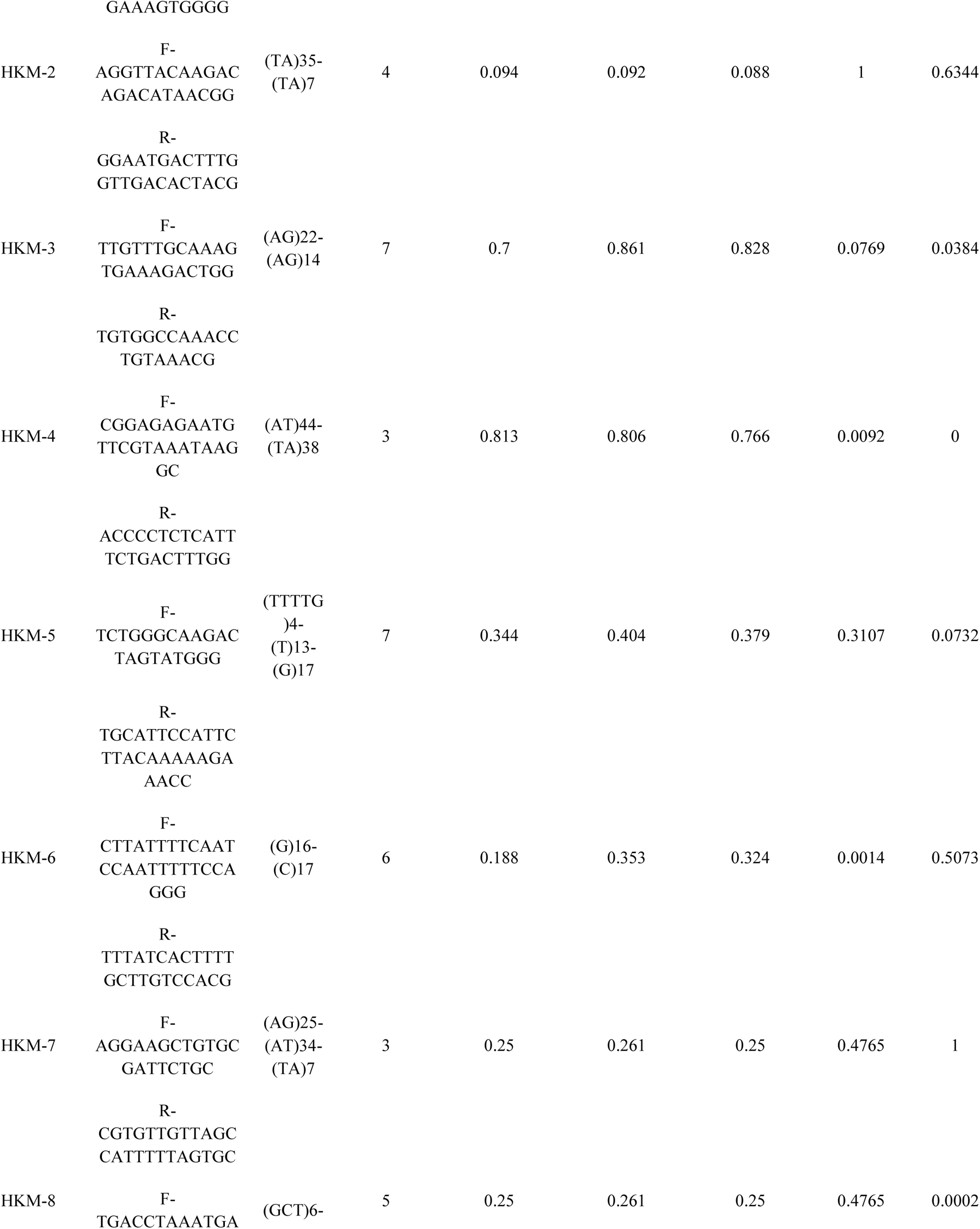
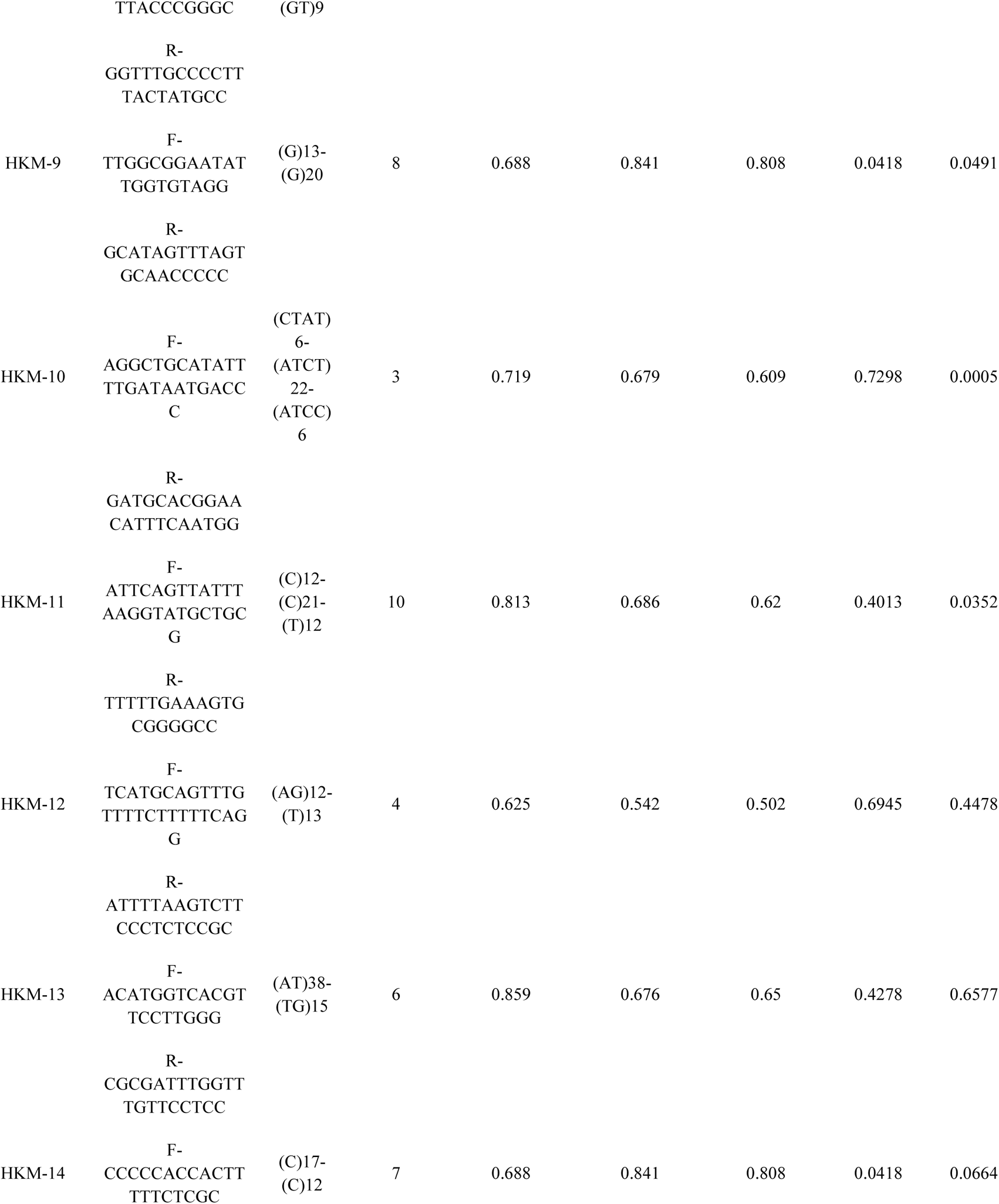
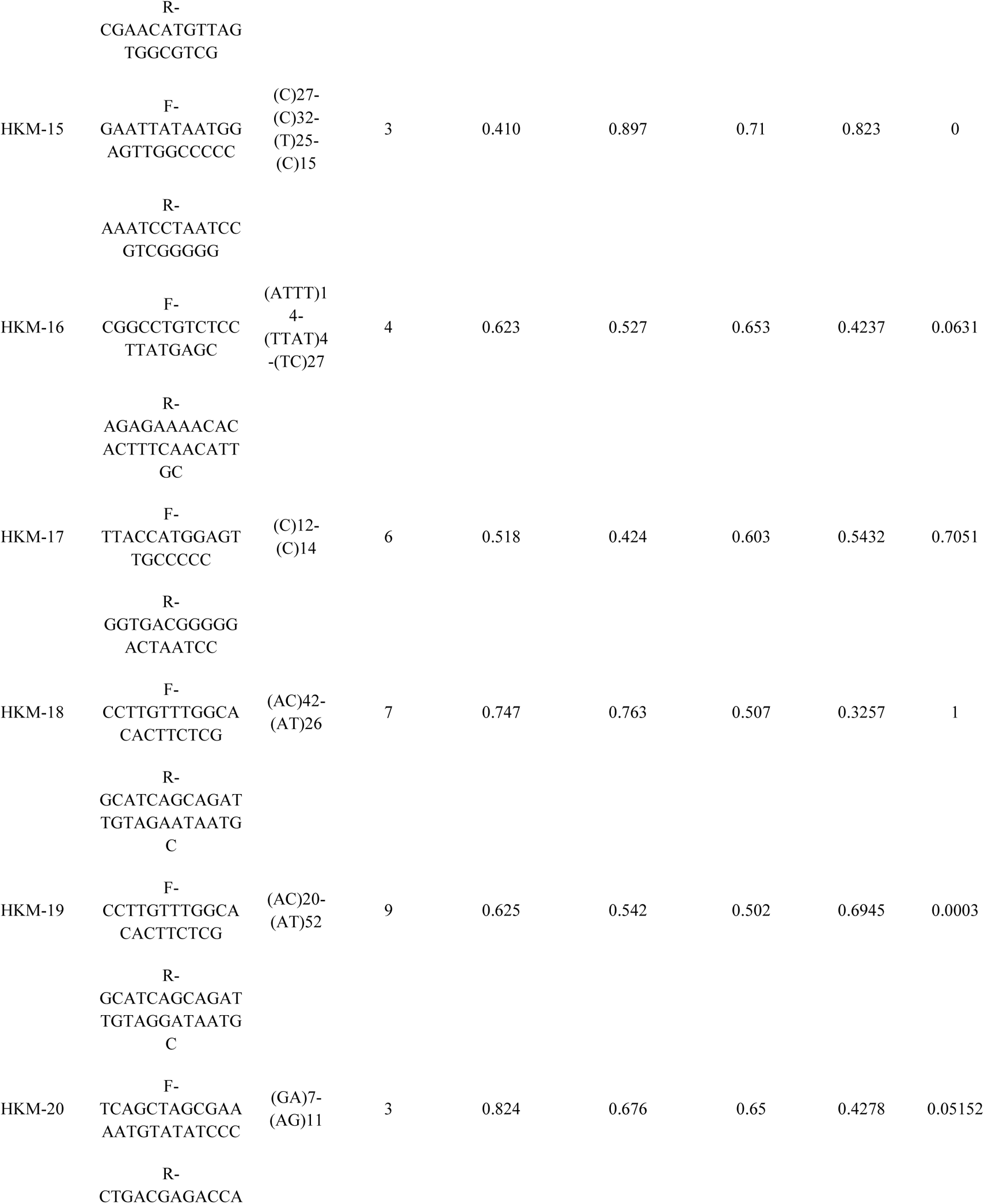
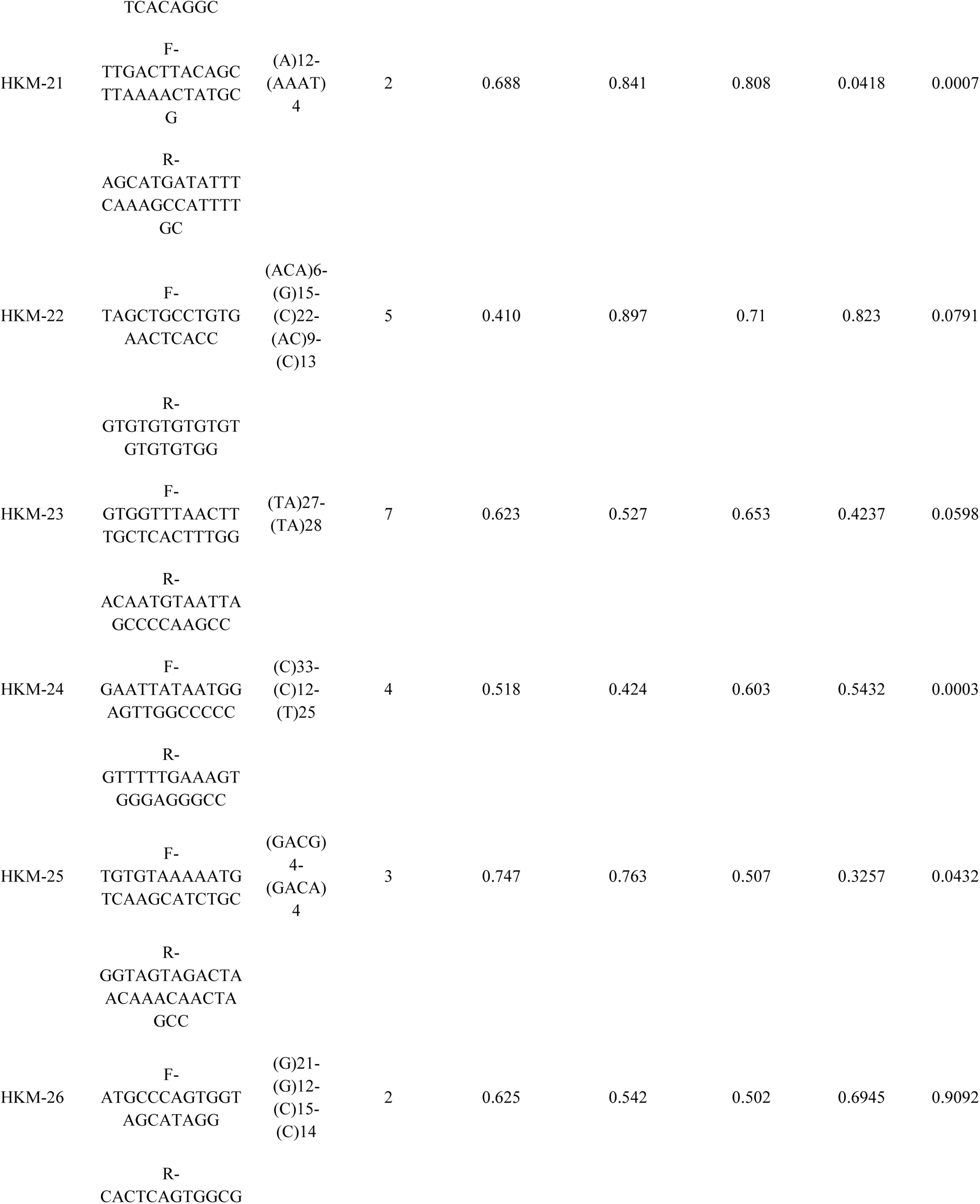
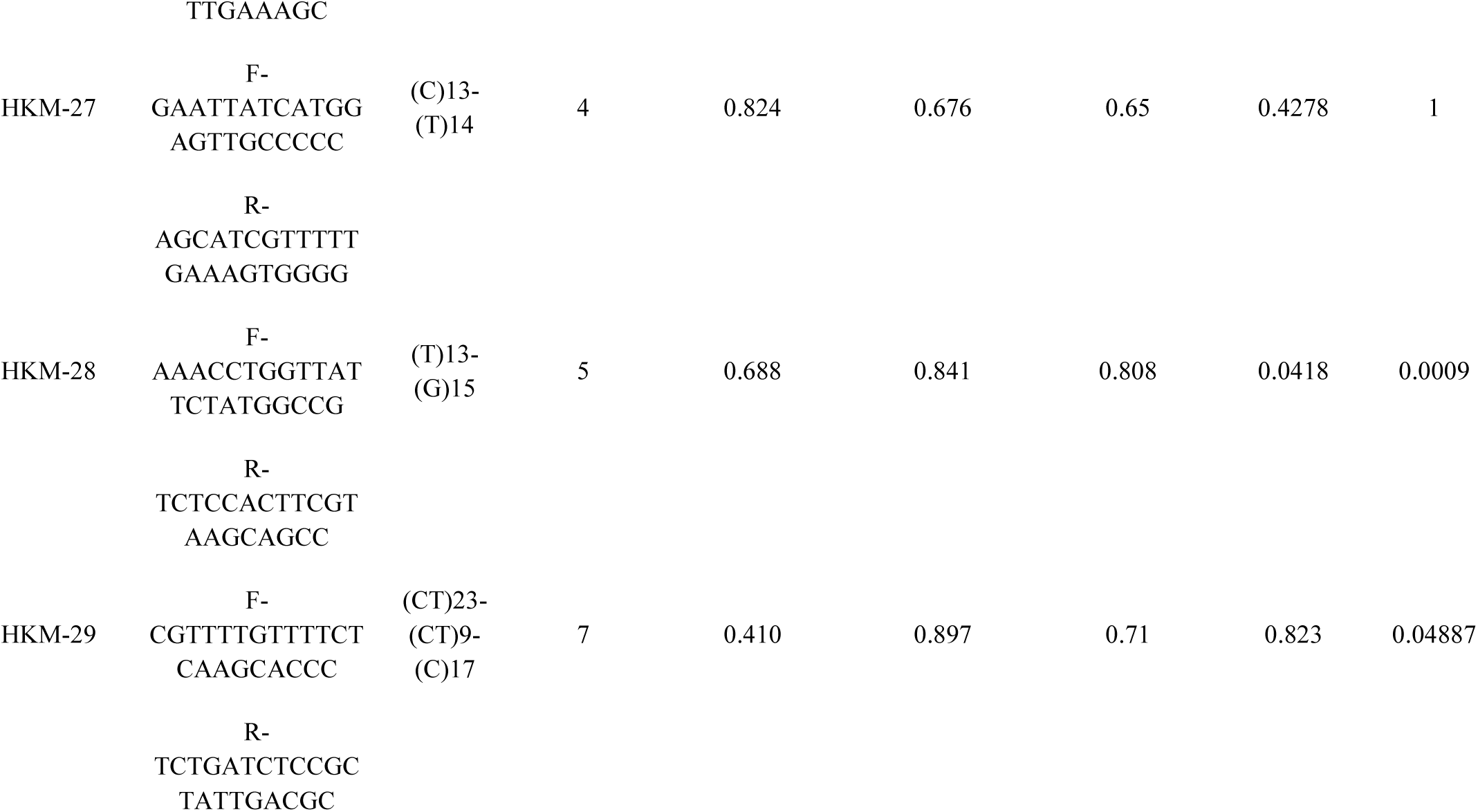
cSSR loci development followed by the diversity analysis within the wild population.

**Table 4:**
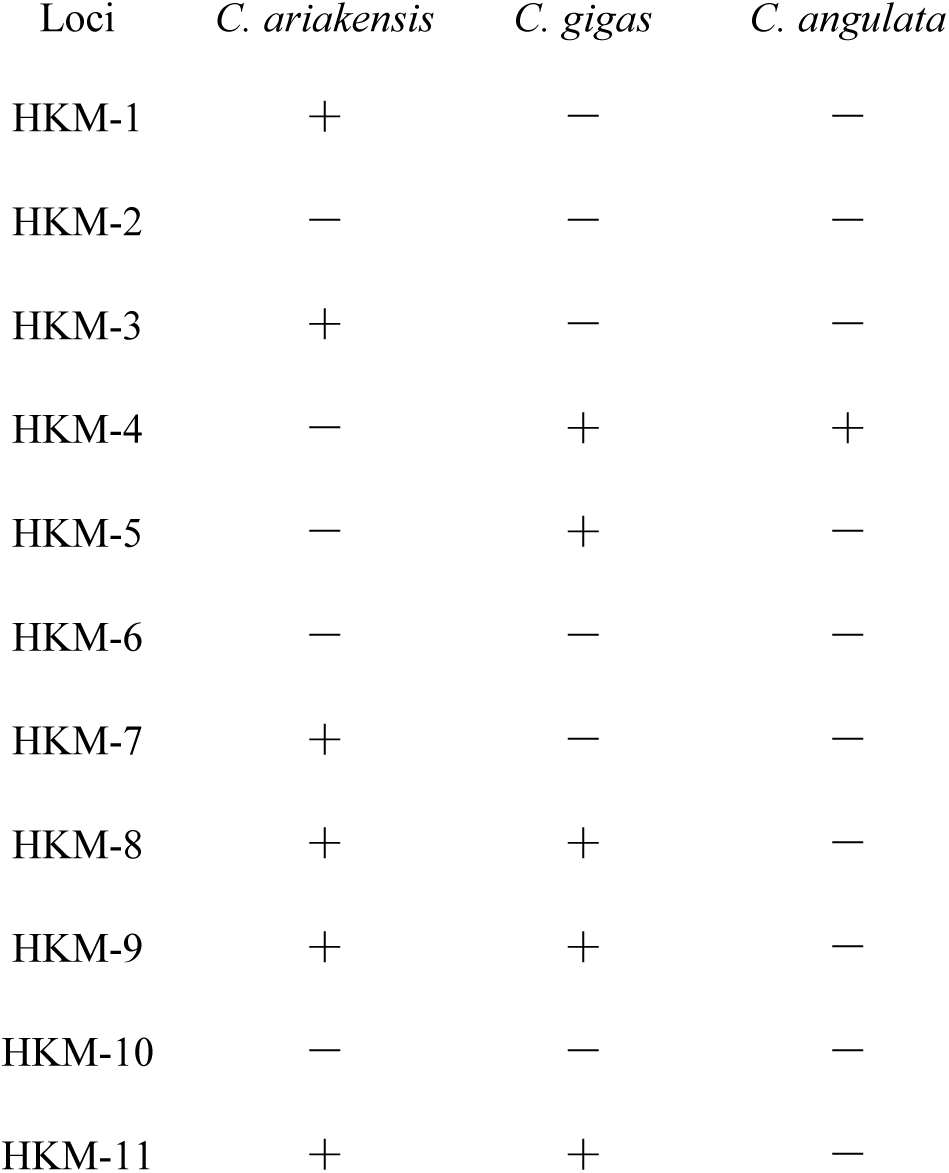
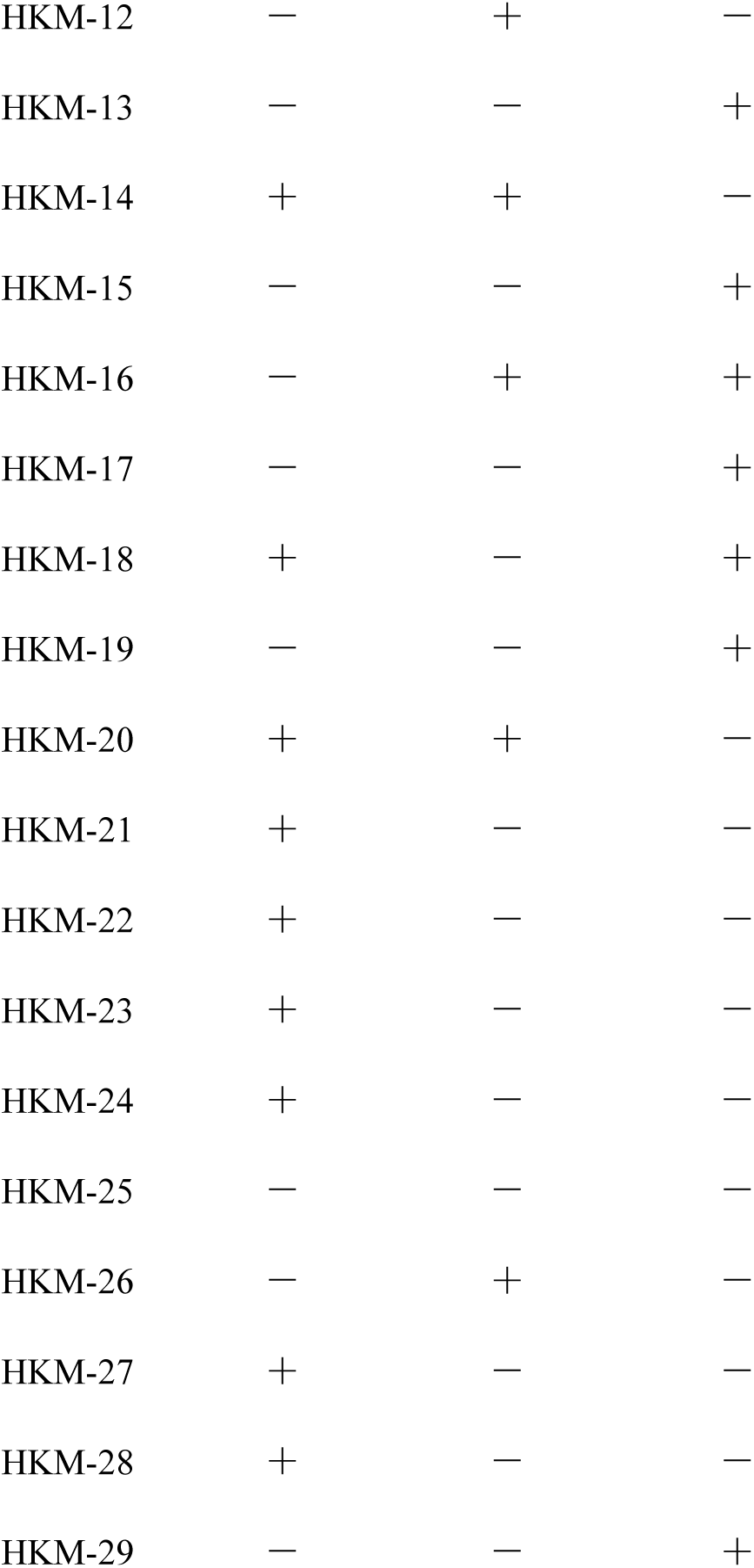
Cross-species amplification of developed loci.

## 4. Discussion

### 4.1. Genome-wide distribution pattern of SSR, motif complexity, and its characteristics

The genome of Crassostrea enriched with SSRs, accomplished 0.76–0.88% of the entire genome sequences which was higher compared with bovine (0.48%) (Qi, Jiang et al. 2015), Chickens (0.49%) (Huang, Zhang et al. 2016), the giant panda (0.64%), (Huang, Li et al. 2015), birds (0.13–0.49%) (Huang, Li et al. 2016), but lower than humans (3%) (Subramanian, Mishra et al. 2003). This variation might be raised due to differences in parameters, algorithms, and software for searching microsatellites or the rate of microsatellite evolution within species (Sharma, Grover et al. 2007). However, this range of SSR occupancy was noticed within the genome of two closely related species such as humans and chimpanzees (Kelkar, Tyekucheva et al. 2008), and between different species of Drosophila (Ross, Dyer et al. 2003). Thus, our result coincides with the other study and suggests that SSR occupancy difference between species might be common phenomena along the taxa. The genus Crassostrea microsatellite frequency enriched with mononucleotide microsatellites followed by di, tetra, tri, penta, and hexa nucleotides, similar to giant panda (Huang, Li et al. 2015), humans (Subramanian, Mishra et al. 2003), arabidopsis (Lawson and Zhang 2006) and mushroom (Wang, Chen et al. 2014). However, the deviation was noticed within rodents (Tóth, Gáspári et al. 2000), where dinucleotide and trinucleotide microsatellites were the most prevalent SSR category. We observed that the incidence of tetranucleotide microsatellites was higher compared with trinucleotide as they engaged to form triplet codes for gene expression. Hexanucleotide microsatellites appeared significantly underrepresented in Crassostrea like other organisms, such as goats (Qi, Jiang et al. 2015) and wild yak (Ma 2015).

In our study, the major repeat motifs were (C/G)n, (AC)n, (AT)n, (AG)n, (AAT)n, (AAC)n, (AAAT)n, (AAAC)n, (AAAG)n, (AAGG)n, (AGAT)n and (AAAAC)n shared by the five Crassostrea species. A maximum of these motifs were enriched with AT compared with GC content which is consistent with many eukaryotes, such as humans (Subramanian, Mishra et al. 2003), and bovine (Qi, Jiang et al. 2015). The following theories may be responsible for the richness AT content in microsatellites: (1) the methylation of thymine (T) (Schorderet and Gartler 1992); (2) AT-rich SSRs may lower the annealing temperature followed by increasing the replication slippage rate of DNA (Lujan, Clausen et al. 2014). Interestingly, mono nucleotide repeats were enriched with (G/C)n, consistent with nematode genomes (Castagnone-Sereno, Danchin et al. 2010). The GC content regulates gene expression, recombination, repetitive element variation and distribution, and evolution of the genome (Du, Zhang et al. 2018). Therefore, it will be interesting to decipher the correlation of recombination frequency with high GC content mononucleotide microsatellites of Crassostrea in future research.

### 4.2. Microsatellite distribution characteristics in each chromosome

We observed that the chromosome size of all the analyzed species of the present study had a positive correlation with the incidence of SSR and cSSRs, but no significant influence on GC content, which was similar to previous results obtained from other eukaryotes (Qi, Jiang et al. 2015, Ding, Wang et al. 2017). Furthermore, we noticed that the chromosome length had no significant correlation with the majority of oyster species’ RA and RD as well as cRA and cRD except *C. hongkongensis.* A negative association between genome length and microsatellite density has been noticed in insects (Ding, Wang et al. 2017), whereas a non-significant association has been noticed in birds (Huang, Li et al. 2016), or primates (Xu, Li et al. 2018). As oysters deal with very adverse environments, one can assume that diverse repeat motifs will introduce various mutations to enrich their evolutionary potential to handle variable environmental conditions.

### 4.3. Compound microsatellite characteristics within the genus Crassostrea

The cSSR% of the present study ranged from 14.99-16.75%, which is higher than revealed in humans, composed of 11% compound microsatellite (Kofler, Schlötterer et al. 2008). However, the possible mechanism that drives to increase of the compound microastellite is still unknown (Kashi and King 2006). The maximum cSSR of the present study is composed of two SSRs. The increase in cSSR complexity reduced the number of cSSRs. The selection pressure might be responsible for an organized frame shift of the respective repeat motif to gain or loss of gene function for the adaptation (Ellegren 2004). The most influential parameter on the occurrence of cSSR was the utmost distance within two adjacent SSR (dMax). As the number of cSSRs increased in a linear manner with increased dMax, we speculate random distribution of cSSR throughout the genome (Kofler, Schlötterer et al. 2008). We observed that the chromosome length of all the oyster species had a significant positive influence on the incidence of cSSRs. Furthermore, the number of cSSRs had a significant correlation with the GC content in *C. gigas* (*R^2^*=0.53; *P=*0.01), (*R^2^*=0.98; *P=*0.008), and *C. ariakensis* (*R^2^*=0.41; *P=*0.04) except *C. virginica* (*R^2^*=0.12; *P=*0.27) and *C. angulata* (*R^2^*=0.0001; *P=*0.09). The cSSR% was significantly correlated with the GC content in all five species, That result added GC along with other three parameters such as ‘species’, ‘chromosome’, and ‘SSR-density’ govern compound microsatellite density. ‘Illegitimate’ recombination within self-complementary microsatellite [CTG]13 [CAG]67 formed the cSSR (Liu, Chen et al. 2010). As our analysis did not notice any such motif, we can assume that recombination is not providing the driving force to form cSSR. Duplicated motif such as (TC)-X-(TC), (AG)-X-(AG), (GATA)-X-(GATA) were prevalent within cSSR. We can speculate that the replication slippage within the microsatellite during replication leads to mutation followed by imperfection to form cSSR (Harr, Zangerl et al. 2000, Dieringer and Schlötterer 2003).

### 4.4. cSSR marker development followed by diversity analysis and cross-species amplification

Although a number of studies represented the development and application of microsatellite markers in Crassostrea, the cSSR has been less explored. Many reports suggest that cSSR is more polymorphic compared to simple SSR (Bull, Pabón-Peña et al. 1999, Lian, Wadud et al. 2006). Thus we targeted the cSSR identified through database mining followed by its diversity estimation and cross-species amplification. The PIC rate of the 29 cSSR loci varied from 0.088-0.828 with the average was 0.591 **(Table 3)**. The Hardy-Weinberg equilibrium (*HWE*) resulted in 22 of the 29 loci followed with the equilibrium, and the rest seven deviated due to heterozygote deficiency within the wild population. Additionally, the presence of inbreeding might be responsible for lowering the heterozygosity. The cross-species amplification results were higher compared with the previous study (Yu, Wang et al. 2010). The higher amplification obtained during this study indicates the flanking sequences of the loci are conserved between these species, and thus, could be useful to decipher the genetic diversity among the species. Mutation within the flanking region leads to mismatch annealing of primer and causes the failure in amplification (Lothe, Andersen et al. 1995). The cSSR loci developed during this study can be useful to do the brood stock analysis of *C hongkongensis* as well as other related species followed by other applications such as marker-assisted selection for sustainable aquaculture production.

## 5. Conclusion

In conclusion, a large number of microsatellite markers were observed throughout the Crassostrea genus. However, we observed an ample amount of similarity as well as noticeable differences between the SSR characteristics within the genus. The similar characteristics explain the conserved nature of microsatellites at the genus level and the difference explained the diversity among the species. However, this needs further in-depth analysis like the “ birth and death model” of microsatellites in Crassostrea to decipher its role. Crassostrea is considered one of the prime members of sustainable aquaculture production, but its speciation as well as its genetic improvement act as major challenges. Thus microsatellite developed during this study can be utilized to an insight into the evolution of the species and the selection of efficient brood stock to carry out genetic improvement programs of the respective species.

